# Distinct lipid profile, low-level inflammation and increased antioxidant defense as a signature in HIV-1 elite control status

**DOI:** 10.1101/2020.07.02.183756

**Authors:** Maike Sperk, Flora Mikaeloff, Sara Svensson-Akusjärvi, Sivasankaran Munusamy Ponnan, Piotr Nowak, Anders Sönnerborg, Ujjwal Neogi

## Abstract

HIV-1 elite controllers (EC) are a rare but heterogeneous group of HIV-1-infected individuals who are able to suppress viral replication in the absence of antiretroviral therapy. The mechanisms of how EC achieve undetectable viral loads remain unclear. This study aimed to investigate host plasma metabolomics and targeted plasma proteomics in a Swedish HIV-1 cohort including EC and treatment-naïve viremic progressors (VP) as well as HIV-negative individuals (HC) to get insights into mechanisms governing EC status. Metabolites belonging to antioxidant defense had higher levels in EC relative to VP while inflammation markers were increased in VP compared to EC. Only four plasma proteins (CCL4, CCL7, CCL20, and NOS3) were increased in EC compared to HC and CCL20/CCR6 axis can play important role. Our study suggests that low-level inflammation and oxidative stress at physiological levels could be important factors contributing to control of viral replication.

**One Sentence Summary:** HIV-1 elite controllers had a distinct lipid profile, reduced inflammation, and increased antioxidant defense that might contribute to elite control status.

## Introduction

HIV-1 elite controllers (EC) are a heterogeneous group among HIV-1-infected individuals who can suppress viremia in the absence of antiretroviral therapy (ART). Although specific definitions for EC vary, it is agreed that EC remains low levels of HIV-RNA and physiological levels of CD4^+^ T cell counts without showing any clinical symptoms of HIV-1 infection for a prolonged period. They are a rare patient subset (<1% of HIV-1-infected individuals) but of great interest in HIV-1 research since they might hold a key for developing HIV-1 cure or vaccines(*1*). It is, however, unclear which mechanisms lead to viral suppression in EC.

It is thought that the major part of viral control can be attributed to host factors rather than viral factors (e.g., infection with a defective or attenuated viral strain). Several small studies have indicated that host genetic factors such as human leukocyte antigen (HLA), for example, HLA-B*57:01, HLA -B*27:05, HLA-B*52, or HLA-A*25 and CCR5 Δ32, can play protective roles in HIV-1 infection(*2*), but larger multi-cohort studies failed to prove this hypothesis. Our earlier proteo-transcriptomic study reported that multiple immune pathways can play a synergistic role in controlling the viral replication in EC and that this group is heterogeneous with distinct properties(*3*).

The progressive development of high-throughput technologies makes it possible to collect and analyze large amounts of genetic and molecular data simultaneously. In this study, we aimed to characterize the metabolic signature of the elite control status in HIV-1-positive individuals and elucidate the mechanism by integrating targeted proteomics and immunological assays from a well-defined EC cohort. We observed EC phenotype-specific lipid profiles that are linked with antioxidant defense and inflammation; together, they might be important factors contributing to control of viral replication.

## Results

### General findings

We used three age-, gender- and body mass index (BMI)-matched cohorts, EC (n=14), viremic progressor (n=16; VP) and HIV-negative controls (n=12; HC) and performed plasma metabolomics analysis using four different ultra-high-performance liquid chromatography and mass spectrometry (UHPLC-MS/MS) methods (Fig. 1A). A total of 950 biochemicals were identified. The majority of them belonged to the class of lipids (49%), followed by amino acids (21%) (Fig. 1B). Group-wise comparison of detected metabolites and percentage of each biochemical class involved is shown in Fig. 1C-E. In the unsupervised principal component analysis (PCA), all samples, regardless of study group, clustered together except for one EC sample (EC06), which was separated from the others (Fig. S1). This sample EC06 was statistically classified as an outlier that could be caused by technical errors and was therefore excluded from further analyses of the metabolomics data. For the remaining 41 samples (EC with n=13, HC with n=12, VP with n=16), a group effect was seen for 294 of the detected biochemicals and similar amounts of altered biochemicals were found when comparing EC vs. VP (236) and VP vs. HC (256). We observed that biochemicals were often decreased in VP compared to the other two groups: of the molecules that were altered compared to EC 86.7% were reduced in VP and of those that were changed compared to HC 87.3% were lower in VP. This indicates a strong metabolic differentiation of VP from the other two groups. The segregation of EC and VP as well as of HC and VP is depicted in volcano plots (Fig. S2, A and B). EC and HC showed only a modest separation (Fig. S2C). Interestingly, HC and EC clusters were most segregated in partial least squares-discriminant analyses (PLS-DA) analysis, whereas VP clustered somewhere between them (Fig. 1F).

**Fig. 1.**
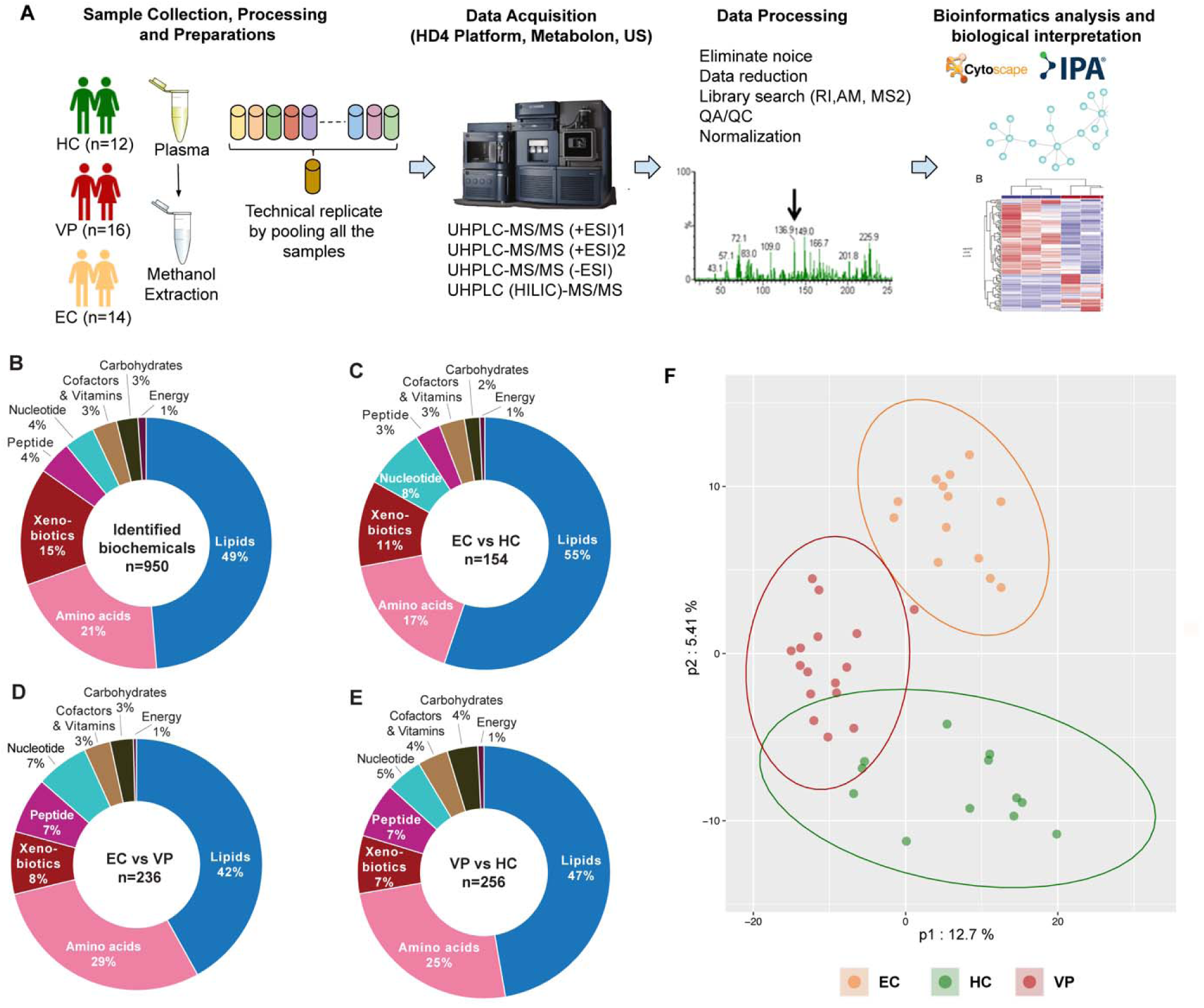
Study design and general findings. (**A**) Study design. Plasma samples of 12 HIV-negative individuals (HC), 16 HIV-1-infected treatment-naïve individuals with viremic progression (VP), and 14 HIV-1 Elite controller (EC) were prepared and analyzed by ultra-high-performance liquid chromatography-tandem mass spectrometry (UHLC-MS/MS). Raw data processing, biochemical identification, quality control, and data normalization was performed according to Metabolon pipeline. Results were analysed by appropriate bioinformatical and statistical methods *in house*. (**B-E**) Doughnut chart with proportions of each super pathway (B) based on the total number of detected and identified metabolites in all groups (950 metabolites) (C) based on the number of metabolites with significantly different levels in EC and HC (p<0.05, 154 metabolites) (D) based on the number of metabolites with statistically significantly different levels in EC and VP (p<0.05, 236 metabolites) € based on the number of metabolites with statistically significantly different levels in VP and HC (p<0.05, 256 metabolites). (**F**) Partial least squares discriminant analysis (PLS-DA) of the 950 in all samples detected and identified metabolites revealing distinct clustering of the three study groups EC (orange), VP (red), and HC (green).

### Components involved in lipid metabolism contribute most to group segregation

To rank the biochemicals according to how important they are for group separation, we used random forest (RF). The top 30 biochemicals revealed by this method are presented and subsumed under their biochemical class (**Error! Reference source not found**.A). A predictive accuracy above 50% implies hereby that results are not random chance. When comparing all three groups, the high predictive accuracy of 90.2% was obtained with biochemicals involved in lipid and amino acid metabolism (11 and 8 out of 30 metabolites respectively) dominating the top-ranked intermediates followed by nucleotide metabolism (6 out of 30 metabolites, **Error! Reference source not found**.A). An even higher predictive accuracy was given for VP vs. EC samples (96.6%); whereas 85.7% was seen for VP vs. HC and 100% for EC vs. HC (Fig. S3). For all of them, key differences were suggested in lipid and amino acid metabolism.

Supervised clustering analyses were used for the top 30 ranked metabolites that were revealed by RF. Samples of the EC and VP group dispersed moderately demonstrating a low metabolic variation within these groups. We observed the segregation of EC and VP with only one VP sample clustering within EC. The clustering was consistent in hierarchical clustering analysis (HCA) (Fig. 2B) as well as in PCA (Fig. 2C) and indicates a robust metabolic differentiation between EC and VP. HC samples were in between those of EC and VP showing a modest separation of EC and HC (Fig. 2, B and C). However, four of the 12 HC samples clustered together with VP implying some differences within this sample group. As already seen in volcano plots belonging to unsupervised analyses, supervised HCA also revealed a decrease of metabolites in VP compared to EC and HC (more than two-thirds of the top 30 metabolites, colored blue in heatmap, Fig. 2B).

**Fig. 2.**
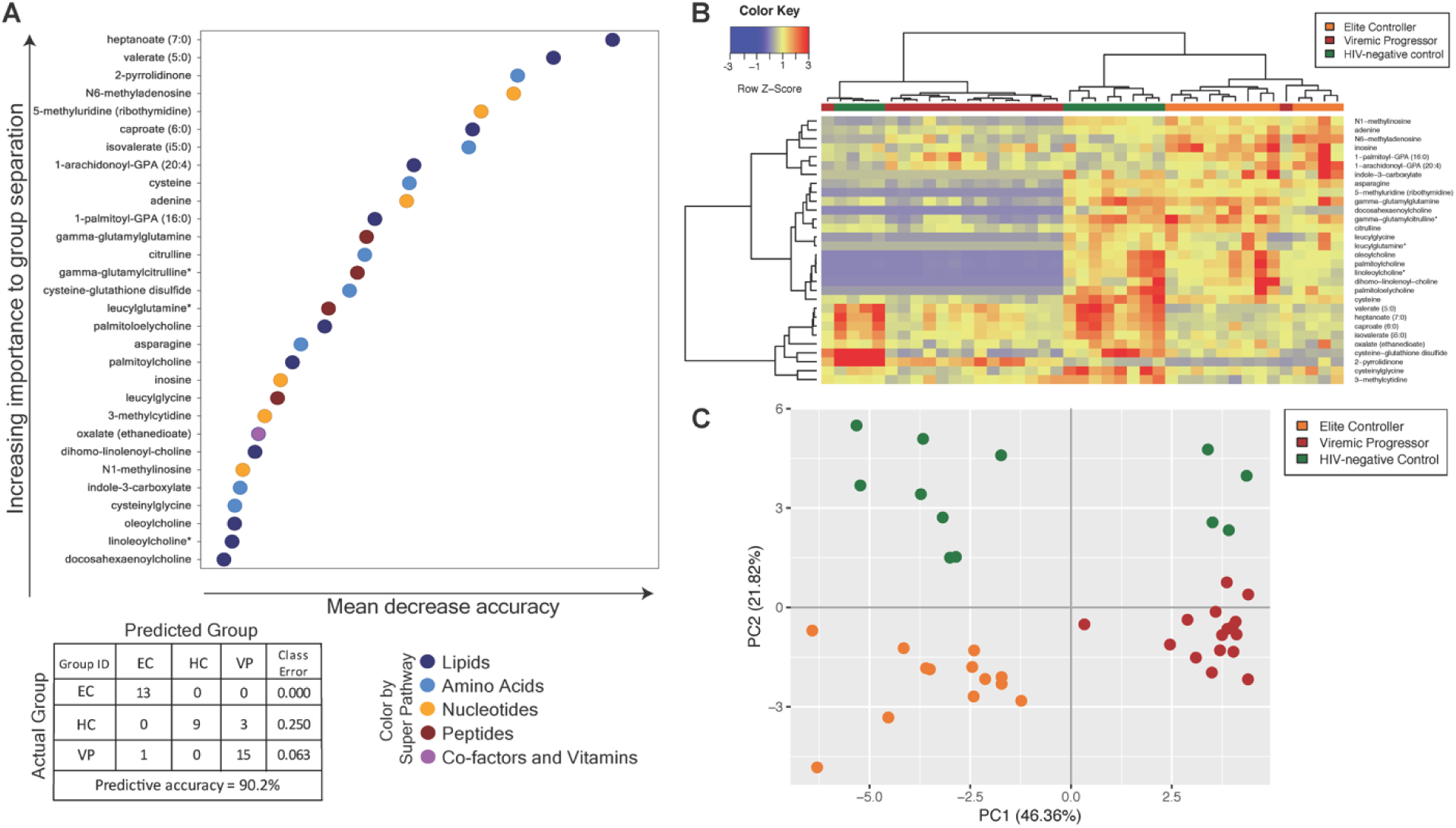
Random forest and supervised analyses of metabolites. (**A**) Random forest (RF) analyses showing the top 30 metabolites that contribute to group separation and the super pathways, the metabolites belong to (see color-coded legend). Biochemicals involved in lipid and amino acid metabolism dominate the top-ranked intermediates. The table represents predictive accuracy, predicted and actual grouping of the samples. (**B**) Supervised hierarchical clustering analyses based on the top 30 metabolites using the Pearson method with Ward algorithm. (**C**) Supervised principal component analyses (PCA) based on the top 30 metabolites.

### Unique acylcholines profile in EC

Pathway mapping of all detected biochemicals reflects the decrease of metabolites in VP compared to EC and HC (Fig. S4). Since most metabolites detected belong to lipids, it is not surprising that most differences between groups were seen for these biomolecules. Concordantly with the general findings, VP had lower lipid levels compared to EC and HC, including for example several metabolites belonging to polyunsaturated fatty acids, lysophospholipids, phospholipid metabolism, or lysoplasmalogen. In contrast, diacylglycerol levels were highest in VP and unchanged in EC compared to HC. No one-sided trend was observed between EC and HC though, not in the metabolic network as a whole and neither for lipids (Fig. 3A). Instead, some lipid subclasses (e.g. lysophospholipds, phospholipid metabolism, as well as primary and secondary bile acid metabolism) were increased, and others (e.g. acylcarnitines, sphingomyelins, phosphatidylcholines, and ceramides) were decreased in EC. An increase or decrease was usually consistent within one subpathway (Fig. 3A).

**Fig. 3.**
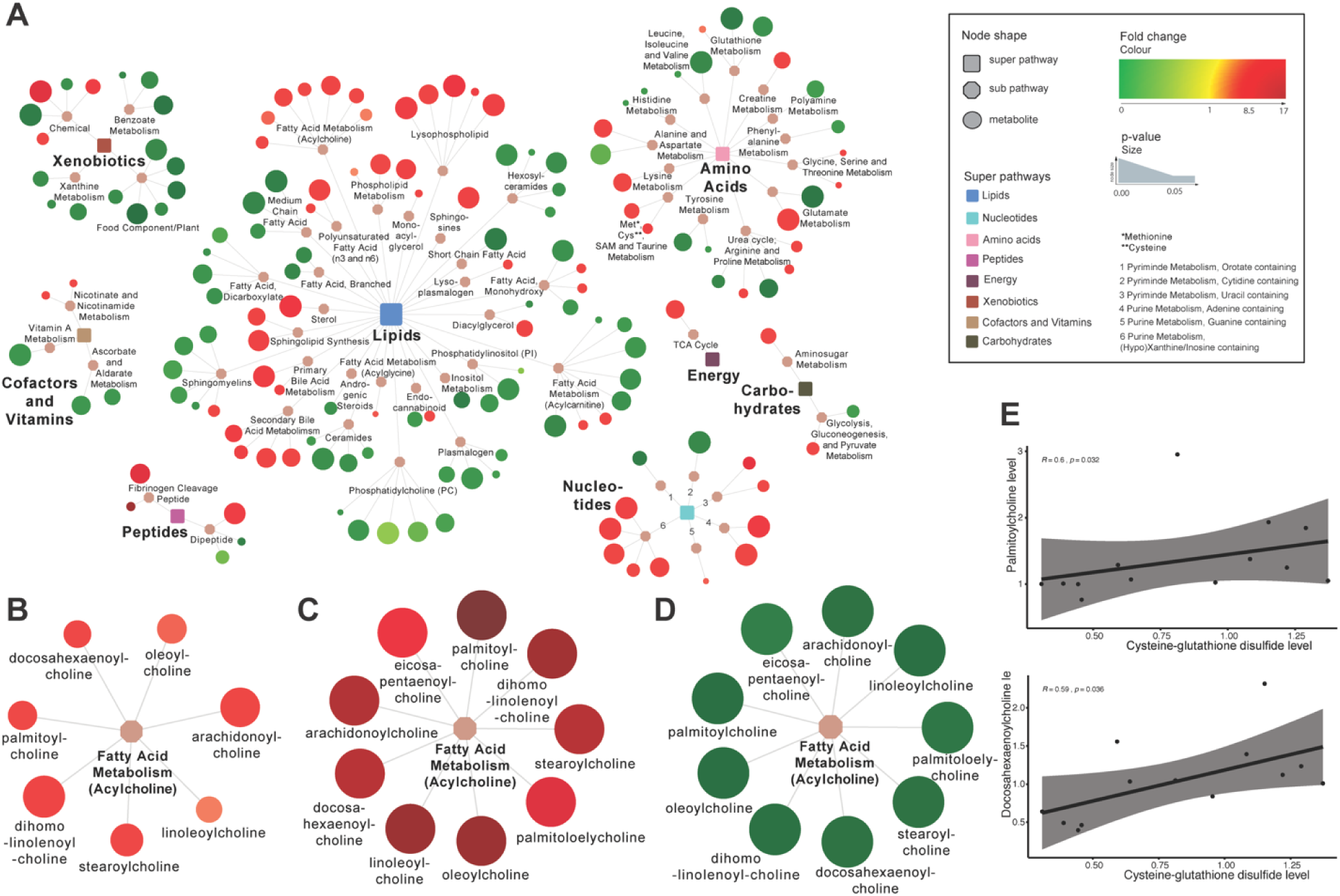
Network analyses. (**A**) Network analyses of the metabolites that were significantly different in EC vs HC (154 metabolites). Rectangular nod shapes represent the eight super pathways that are shown in different colors according to legend. Octagonal nod shapes are used for sub pathways belonging to the eight super pathways. Circular nod shapes show the single metabolites, where red indicates increased levels and green indicates decreased levels in EC relative to HC. Size of the circles’ indicate p-value: the bigger the size the lower the p-value. Lines connect each metabolite to its respective subpathway and subpathway to their respective super pathways. (**B-D**) Network showing the sub pathway *Fatty acid metabolism acylcholines* and metabolites belonging to it with red indicating increased levels and green indicating decreased levels. Size of the circles picture p-value (B) EC vs. HC (C) EC vs VP (D) VP vs HC (**E**) Spearman correlation analyses between metabolites were performed for EC samples and revealed a moderate correlation between levels of cysteine-glutathione disulfide and of palmitoylcholine (R=0.6, p-value=0.032) as well as levels of cysteine-glutathione disulfide and of docosahexaenoylcholine (R=0.59, p=0.036). Respective scatter plots are presented.

We further observed that lipids being part of acylcholines were reduced in VP and elevated in EC compared to HC. Every acylcholine detected was significantly and markedly decreased in VP subjects relative to HC. Interestingly, acylcholines were not decreased, instead, most of them were even increased in EC relative to HC, leading to markedly elevated acylcholines in EC relative to VP (3.5 <fold change< 17.5, Fig. 3B-D). Of note, choline levels were also elevated in EC compared to HC (p=0.044) and VP (p<0.001).

Interestingly, levels of two of the detected acylcholines (palmitoylcholine and docosahexaenoylcholine) correlated with the level of cysteine-glutathione disulfide that is involved in antioxidant defense (R=0.6 and p=0.032 for palmitoylcholine and R=0.59 and p=0.036 for docosahexaenoylcholine) (Fig. 3E).

### Increased antioxidant defense in EC relative to VP linked to glutathione and one-carbone metabolism

As the choline levels may be linked to changes in oxidative stress and/or antioxidant defense, next we investigated glutathione metabolism linked with transsulfuration pathway and one-carbone metabolism as the cellular tripeptide glutathione is a key cellular antioxidant compound regulating redox homeostasis (Fig. 4A). Relative to VP, EC showed increases in the methylation reaction product S-adenosylhomocysteine (SAH) (p<0.001), the glutathione precursors cysteine (p<0.001) and glycine (p=0.004), the cysteine-derived antioxidants hypotaurine (p<0.001) and taurine (p<0.001), as well as the glutathione cycle intermediates cysteinylglycine (p=0.004), 5-oxoproline (p<0.001), glutamate (p<0.001) and several gamma-glutamyl amino acids (Fig. 4A, Fig. S5, and Table S1). Together, these data suggest that EC exhibit increased activity of the transsulfuration pathway toward *de novo* glutathione synthesis in addition to increased glutathione recycling compared to VP. Additionally, methionine, which is linked to glutathione synthesis via the transsulfuration pathway, was also increased in EC compared to VP (p=0.003), suggesting the aforementioned increases in glutathione synthesis in EC may also reflect elevated methionine availability. Furthermore, EC showed, relative to VP, increases in many oxidized intermediates including cysteine-glutathione disulfide (p=0.012), cysteinylglycine disulfide (p=0.002), oxidized cysteinylglycine (p=0.04), cystine (p=0.007), methionine sulfoxide (p<0.001), and methionine sulfone (p<0.001) (Supplementary S6a). Looking at EC and HC, significant increase was observed in 5-oxoproline (p<0.046), glutamate (p<0.001), serine (p=0.048) and SAH (p=0.02) in EC (Fig. 4A and Fig. S5). The level of metabolites that are part of amino acids; methionine, cysteine, SAM, taurine and glutathione metabolism and that are linked to above mentioned one-carbon metabolism, transsulfuration, gamma-glutamyl cycle and redox homeostatis is presented in Fig. 4B. While increases in oxidized intermediates could reflect elevated oxidative stress, in the context of elevated glutathione synthesis/recycling and cysteine availability, this signature likely also reflects improved antioxidant defense mechanisms and greater detoxification of reactive oxygen species (ROS) in EC.

**Fig. 4.**
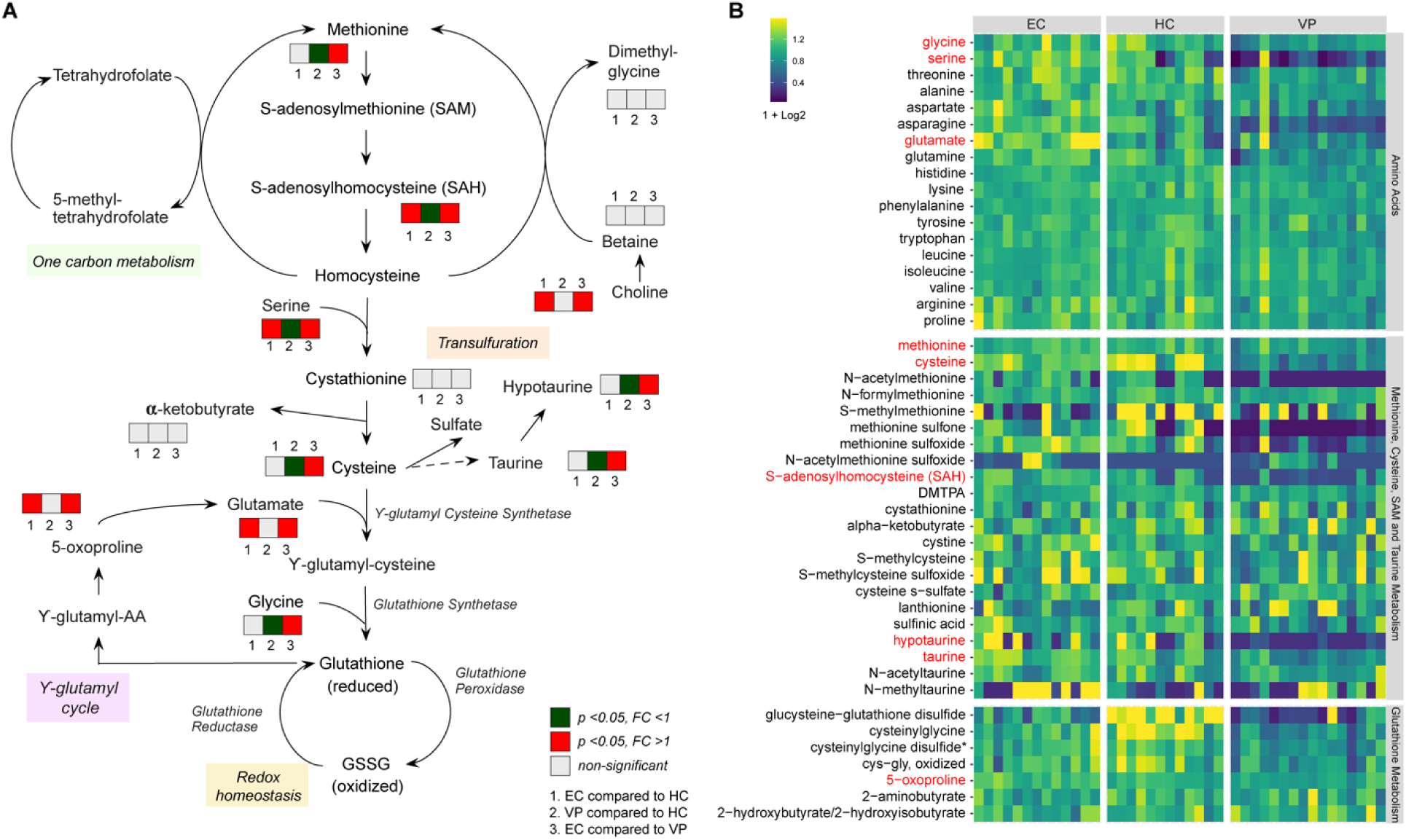
Methionine, Transsulfuration and Glutathione metabolism. (**A**) A schematic presentation of methionine metabolism that is linked to transsulfuration pathway, one-carbon metabolism, and glutathione metabolism. Methionine is transformed into S-adenosylmethionine (SAM) and S-adenosylhomocysteine (SAH) and finally converted into homocysteine, which is also connected to one-carbon metabolism. In the transsulfuration pathway cysteine and homocysteine are interconverted through the intermediate homocysteine. Cysteine can give rise to the antioxidants taurine and hypotaurine and it is also part of *de novo* glutathione synthesis. As glutamate is one of the three peptides glutathione (GSH) constitutes of, there is a link to the gamma-glutamyl cycle. By reducing reactive oxygen species (ROS), GSH is transformed to its oxidized form GSSG and GSSG can be reverted to reduced GSH by the enzyme glutathione reductase. Boxes in neighbourdhood of a metabolite indicate that the respective metabolite was detected and quantified in all our samples. Box 1 shows comparison between EC and HC, box 2 betweenV P and HC and box 3 between EC and VP. A grey-coloured box means non-significant difference in plasma levels, red presents foldchange greater than one (with p-value<0.05) and green foldchange smaller than one (with p-value<0.05). (**B**) Heatmap representing levels of metabolites that are part of amino acids; methionine, cysteine, SAM and taurine metabolism; and glutathione metabolism. Samples are grouped according to study group (EC, HC, VP). Metabolites written in red are part of the scheme shown under (A). Colour depictures increasing log2 levels from blue via green to yellow.

### Inflammation markers in EC relative to HC

As any kind of infection is often accompanied by systemic inflammation, and there is a known link between oxidative stress and inflammation, we were next interested in levels of inflammation markers. We hypothesized that improved antioxidant defense allows EC to keep the inflammation level low despite an earlier study suggested increased inflammation in EC compared to HC(*4*). On a metabolic level, we noticed lower levels of kynurenine and quinolinate together with increased levels of serotonin and tryptophan in EC compared to VP, but no differences between EC and HC. This signature likely reflects an increased metabolism of tryptophan to kynurenine along kynurenine pathway in VP (but not in EC), which is associated with inflammation(*5*) (Fig. 5, A and B). To further support this we used plasma proteomics markers of inflammation in HC and EC and compared levels of selected plasma proteins using two different approaches: re-analysis of data that was obtained by proximity extension assay (PEA) applying Olink immuno-oncology panel, which has partly been published earlier(*3*); together with enzyme-linked immunosorbent assay (ELISA) for detection of C-reactive protein (CRP), and neopterin. In the proteomics data analyses, EC06 was included (EC with n=14, HC and VP as for metabolomics). Out of 92 plasma proteins analyzed by PEA, 81 passed quality control and were used for statistical analyses. Partial least squares-discriminant analysis (PLS-DA) showed group clustering with VP being separated from the other two groups, whereas HC and EC intermingled to some extent (Fig. 5C). This clustering is similar to that of the metabolomics data. Out of the 81 proteins, four were statistically different in EC and HC: CCL4 (p=0.001), CCL7 (p=0.002), CCL20 (p<0.001), and nitric oxide synthase 3 (NOS3, p=0.006) (Fig. 5D). All of them were increased in EC compared to HC. ELISA confirmed elevated CCL20 levels in EC relative to HC (data not shown). Median (IQR) CRP levels in the study groups indicate a trend of elevated CRP plasma levels in EC (1.32 [0.95-2.49]) compared to HC (1.07 [0.69-1.71]), but lower than in VP (2.38 [2.25-3.66]). That was, however, statistically not significant (EC vs HC p=0.2469 and EC vs VP p=0.1469) (Fig. S6A). Neopterin plasma levels were increased in VP, but not in EC compared to HC (Fig. S6B). This data indicates that EC has a rather similar inflammation profile to HC and different from VP.

**Fig. 5.**
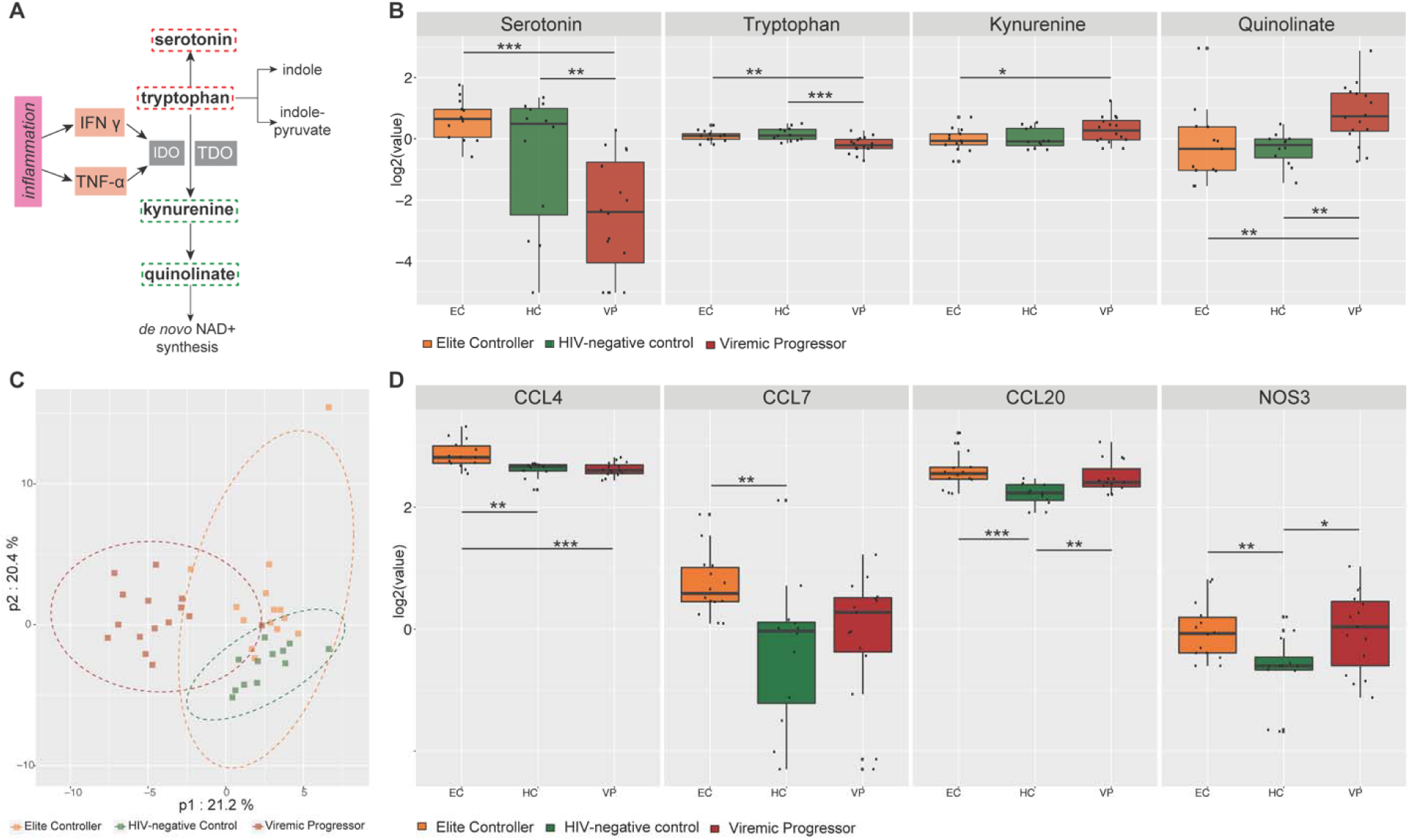
Inflammation markers. (**A**) Tryptophan can be metabolized along several distinct pathways either giving rise to the neurotransmitter serotonin, along the kynurenine pathway that has been associated with inflammation and disease, or to several indole compounds. Kynurenine in liver is generated by tryptophan dioxygenase (TDO) but in extra-hepatic tissues by the enzyme indoleamine 2,3-dioxygenase (IDO), which is well characterized to be induced by the inflammatory cytokines, interferon-gamma (IFNγ) and tumor necrosis factor-alpha (TNF-α). Kynurenine can further be metabolized to quinolinate that is a precursor of nicotinamide adenine dinucleotide (NAD^+^). Metabolites that were detected in samples are marked with rectangular-shaped broken lines around them; red indicates higher levels and green represents lower levels in EC relative to VP. IDO – Indoleamine 2,3-dioxygenase, IFN γ - Interferon gamma, NAD^+^ – nicotinamide adenine dinucleotide, TDO – tryptophan 2,3-dioxygenase, TNF-α - tumor necrosis factor alpha. (**B**) Boxplots of metabolites that belong to kynurenine pathway. Log2 of serotonin, tryptophan, kynurenine, and quinolinate are presented for EC (orange), HC (green), and VP (red). Median values and interquartile ranges are indicated by bars. P-values are determined by Mann-Whitney U test with * indicating p-value<0.05, **p-value<0.01, and ***p-value<0.001. (**C**) Partial least squares discriminant analysis (PLS-DA) including 81 proteins analysed in plasma samples showing clustering of the three study groups EC (orange), HC (green), and VP (red) samples where VP samples cluster separately from HC and EC, whereas HC and EC intermingle to a small extend. (**D**) Boxplots of proteins that had significantly different levels in HC and EC (p<0.05) revealed by Mann-Whitney U test. Log2 of CCL4, CCL7, CCL20, and NOS3 are presented for EC (orange), HC (green), and VP (red). Median values and interquartile ranges are indicated by bars. *p-value<0.05, **p-value<0.01, ***p-value<0.001.

### Decreased CCR6 surface expression in EC

Cytokine-mediated signaling occurs through binding and subsequent activation of cytokines to their specific receptors. Alterations in either cytokine levels, in expression profiles of their specific receptors, or in both could indicate changes in cytokine signaling pathways. Due to elevated plasma levels of the cytokines CCL4, CCL7, and CCL20 in EC relative to HC, we were interested in the surface expression of their respective receptors, CCR2 (ligand CCL7), CCR3 (ligand CCL7), CCR5 (ligand CCL7 and CCL4), and CCR6 (ligand CCL20), on peripheral blood mononuclear cells (PBMCs). Flow cytometry analyses were therefore performed on a subpopulation of gender-matched EC (n=14) and HC (n=8) samples of the study cohort. Antibody panel was chosen to discriminate CD4^+^ T cells (CD3^+^CD4^+^), CD8^+^ T cells (CD3^+^CD8^+^), and monocytes expressing a classical (CD14^+^CD16^-^), non-classical (CD14^-^ CD16^+^), and intermediate (CD14^+^CD16^+^) phenotype. EC had significantly decreased numbers of CD4^+^ T cells and significantly increased counts of CD8^+^ T cells compared to HC (Fig. 6, A and B). No differences were seen in the proportion of monocytes (Fig. 6, A and C). To our surprise, percentages of cells expressing CCL20 receptor CCR6 were reduced in EC in all cell subpopulations (Fig. 6D) except for intermediate monocytes (Fig. S7A). For CD4^+^ and CD8^+^ T cells the reduction was substantial, while only 0.07% of classical monocytes and 0.17% of non-classical monocytes were CCR6^+^ in HC making the difference to EC marginal (CCR6 expression in 0.02% of classical and <0.01% of non-classical monocytes). Furthermore, surface expression levels of CCR6 were reduced on CCR6 expressing CD4^+^ and CD8^+^ T cells in EC compared to HC (Fig. 6E). Small differences were seen regarding frequency and distribution of CCR2 expression (Fig. 6F). Frequencies of CCR2 expressing cells were reduced in CD8^+^ T cells (7.81% vs 17.00%) and in non-classical monocytes (0.16% vs 0.72%) in EC, while CCR2 surface expression levels were decreased on CD4^+^ and CD8^+^ T cells in EC compared to HC (Fig. 6G). Different distribution of CCR2 and CCR6 expression on CD4^+^ and CD8^+^ T cell populations are depicted in t-SNE plots (Fig. 6H). The frequency of CCR3 expressing cells was generally very low in all cell populations, with an upregulation in CD4^+^ T cells (0.58% vs 0.29%) and downregulation in non-classical monocytes (<0.01% vs 0.13%) in EC (Fig. S7A). The frequency of cells expressing CCR5, which is receptor for CCL7 and CCL4, and also a co-receptor for HIV-1 entry, was similar in EC and HC in all examined cell populations (Fig. S7A). Although a trend was seen for higher frequencies of CCR5 expressing CD8^+^ T cells in EC, that reached not statistical significance (p=0.068) (Fig. S7A). No differences were seen for any of the receptors in intermediate monocytes (Fig. S7A). Analysis of co-expression of several receptors revealed significant differences between EC and HC in both CD4^+^ (Fig. 6I) and CD8^+^ (Fig. 6K) T cell subsets that co-express CCR2, CCR5, and CCR6 with lower frequencies in EC compared to HC. Further, amount of CD4^+^ T cells co-expressing CCR5 and CCR6 as well as CCR2 and CCR6 is reduced in EC relative to HC. The clustering of different cell subsets and expression of surface markers is also depicted in t-SNE plots (Fig. S7b). This data indicates that CCR6/CCL20 chemokine axis along with CCR2/CCR5/CCL4/CCL7 signaling, which plays a role in HIV-1 entry and in antiviral immunity (*6, 7*), are modulated in EC. A summary of important findings in our EC cohort is presented in Fig. 7.

**Fig. 6.**
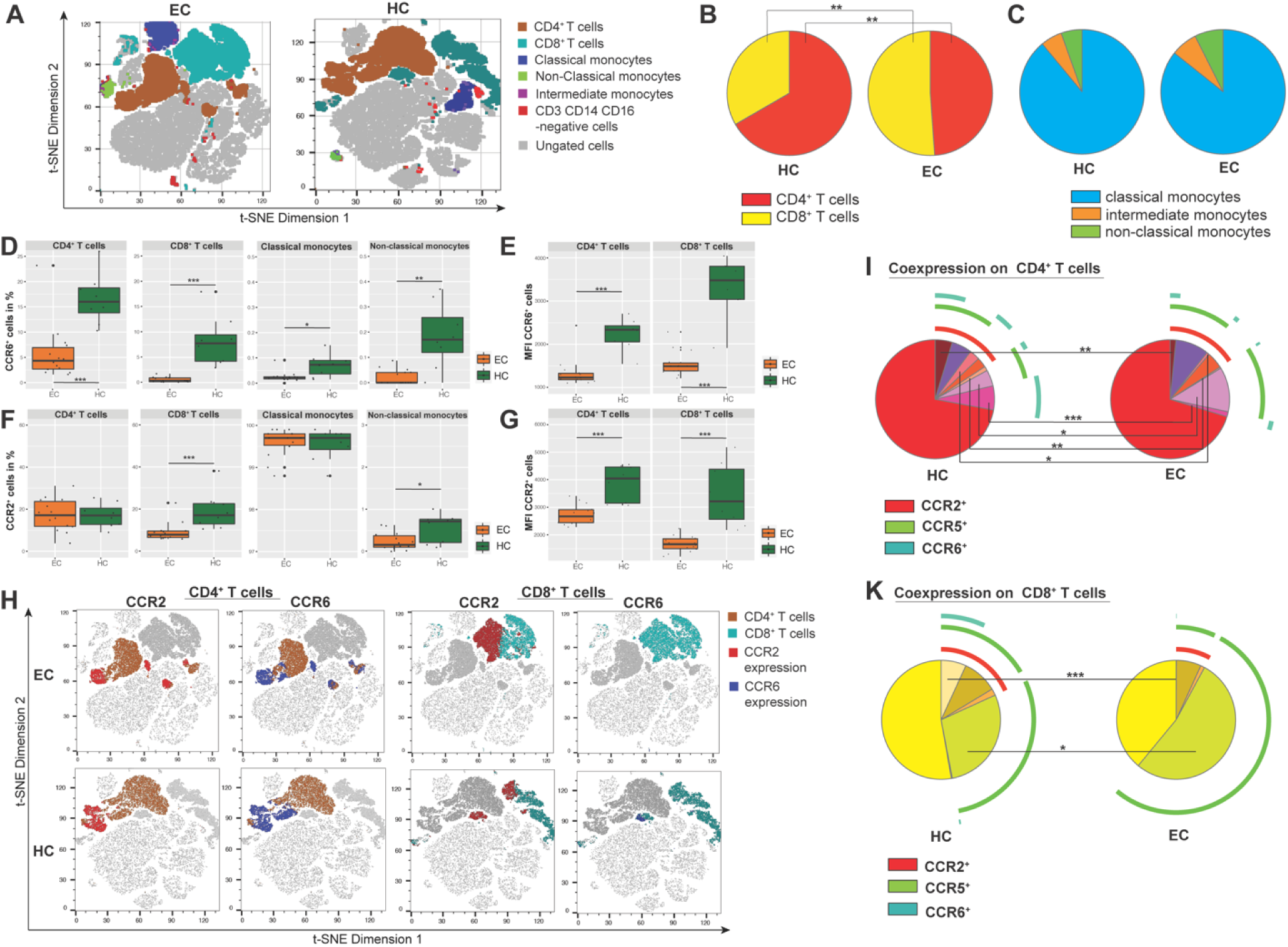
Flow cytometry analyses. (**A**) Differences between EC (left) and HC (right) in overall clustering of PBMCs acquired by flow cytometry. Viable singlet cells of each sample were downsampled to 10,000 and individual downsampled samples of EC and HC respectively were concatenated. t-SNE analysis was performed with 2,000 iterations with perplexity of 20 and learning rate of 1,000. The two plots show t-SNE dimension 1 and t-SNE dimension 2 based on expression of CD4, CD8, CD14, CD16, CCR2, CCR3, CCR5, and CCR6. CD4^+^ T cells are shown in brown, CD8^+^ T cells in light blue, classical monocytes in dark blue, non-classical monocytes in green, intermediate monocytes in purple, cells that are triple-negative for CD3, CD14, and CD16 in red; and ungated cells in grey. (**B**) Pie charts depicture different proportions of CD4^+^ (yellow) and CD8^+^ (red) T cells in HC (left) and EC (right). (**C**) No difference is observed in the proportion of classical (blue), intermediate (orange), and non-classical (green) monocytes between HC (left) and EC (right). Proportions are presented in pie charts. (**D-G**) Boxplots represent expression frequency or median fluorescence intensity (MFI) of selected surface receptors on different cell populations, revealed by flow cytometry analysis, for EC (orange) and HC (green). Median values and interquartile ranges are indicated by bars. P-values are determined by Mann-Whitney U test with * indicating p-value<0.05, **p-value<0.01, and ***p-value<0.001. (D) Expression frequency of CCR6 on CD4^+^ T cells and CD8^+^ T cells as well as on classical and non-classical monocytes. (E) MFI of CCR6 on CD4^+^ T cells and CD8^+^ T cells. (F) Expression frequency of CCR2 on CD4^+^ T cells and CD8^+^ T cells as well as on classical and non-classical monocytes. (G) MFI of CCR2 on CD4^+^ T cells and CD8^+^ T cells. (**H**) Differences between EC (upper row) and HC (lower row) in surface expression of CCR2 and CCR6 on CD4^+^ and on CD8^+^ T cells. t-SNE analysis of downsampled and concatenated samples was performed as described under *a*. Presented t-SNE plots show t-SNE dimension 1 and t-SNE dimension 2. CD4^+^ T cells are highlighted in brown in the two columns on the left and CD8^+^ T cells are shown in light blue in the two columns in the right. Lymphocytes expressing CCR2 are coloured in red and those expressing CCR6 in dark blue. (**I** and **K**) Pie charts depicture co-expression of CCR2, CCR5, and CCR6 on lymphocytes for HC (left) and EC (right). The expression of CCR3 is very low and has therefore been neglected in these illustrations. (I) Co-expression on CD4^+^ T cells (K) Co-expression on CD8^+^ T cells.

**Fig. 7.**
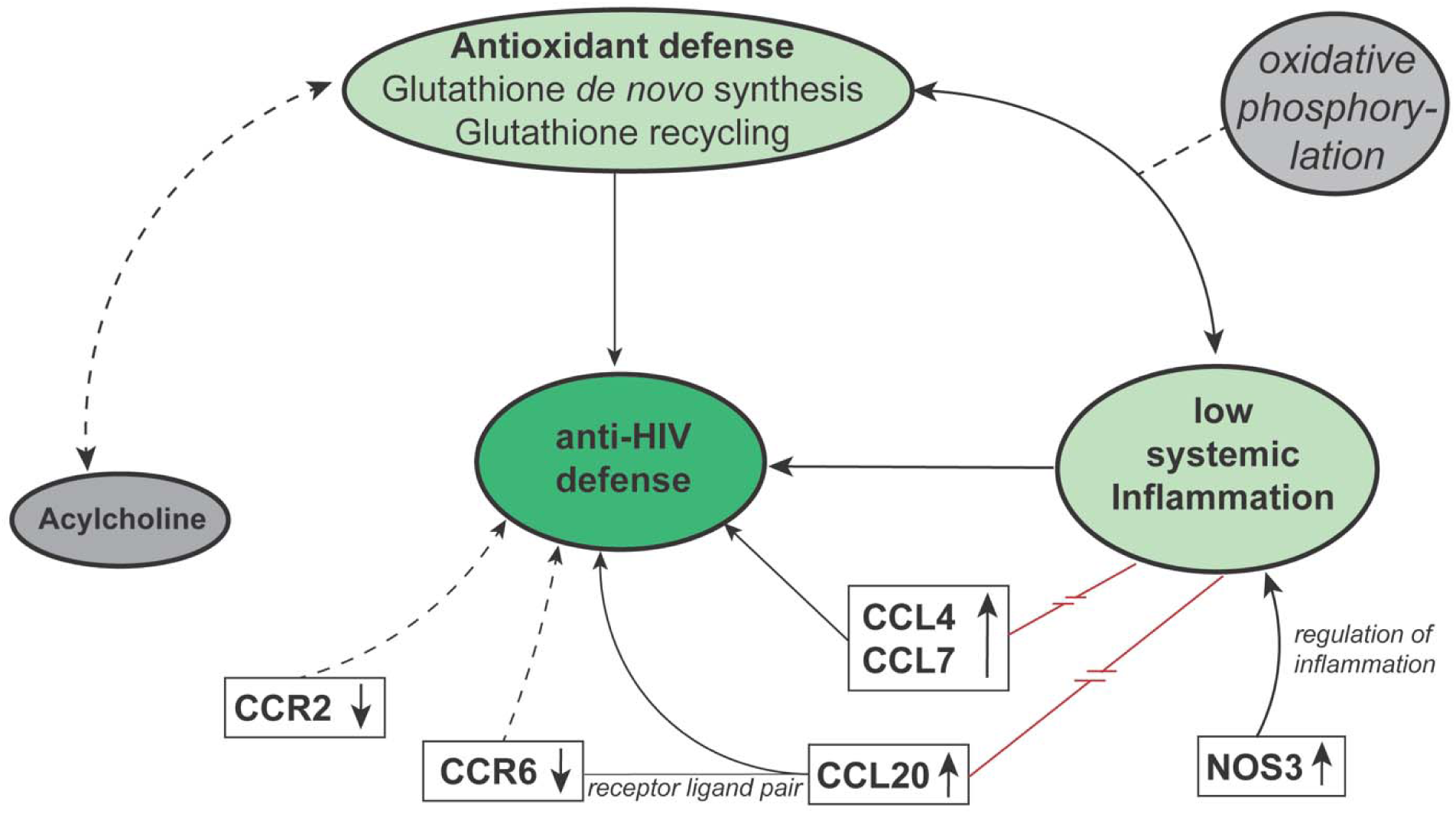
The proposed hypothesis of viral control mechanisms in EC. A summary of our findings that might contribute to viral control. EC showed a physiological state of antioxidant defense and low systemic inflammation with the exception of increased plasma levels of CCL4, CCL7, CCL20, and NOS3. Chemokine receptors CCR2 and CCR6 were downregulated on lymphocytes in EC. Oxidative phosphorylation is hypothesized to be a link between antioxidant defense and inflammation. Acylcholine levels in plasma samples of EC were increased, but biological meaning is of this finding is not known. Arrows depicture known relations between features. Red-coloured line illustrates conflicting findings. Broken lines with arrows represent hypothesized relations between features that need to be investigated in future studies.

## Discussion

Our study indicated a distinct plasma metabolic signature with an increased antioxidant defense that was linked to lower inflammation as a hallmark of EC phenotype. The levels of acylcholins were markedly increased compared to HC and VP, making it a unique feature of EC. These features might be important factors contributing to control of viral replication. Moreover, distinct cytokine-mediated signaling attributed to CCR2, CCR5 and CCR6 can play an essential role in viral tropism and antiviral immunity contributing to the EC phenotype.

Lipid abnormalities have been reported in HIV-1-infected individuals as a result of infection (but also as an effect of ART). By altering intracellular pathways involved in cellular metabolism in general or lipid signaling in particular, an optimal environment can be created for viral replication(*8*). Despite heterogenous in nature, the acylcholines showed marked increases in EC. The biological role of acylcholines is not well understood. Chemically, acylcholines are esters formed from choline and a carboxylic acid and the acylcholines we detected mainly derived from long-chain fatty acids(*9*). Thus, the fatty acids and their salts respectively or choline itself might have a biological function in EC status. One of the acylcholines detected, namely arachidonoylcholine, is known as a cholinergic agonist that binds to and activates cholinergic receptors(*9*). Whether arachidonoylcholine or other acylcholines (besides the classical agonist acetylcholine) and acylcholin receptor signaling play a role in EC viral control needs further investigation. Furthermore, a protective effect of arachidonoylcholine, oleoylcholine, linoleoylcholine, and docosahexanoylcholine against H_2_O_2_-induced cytotoxicity has been discussed(*10*).

Interestingly, acylcholine levels in EC also correlated with levels of metabolites involved in glutathione metabolism. Methionine is a key molecule involved in the regulation of several metabolic processes, such as sulfur metabolism and redox homeostasis. During a process called metoxidation, the sulfur residues in methionine can also be oxidized by ROS into methionine sulfoxide or methionine sulfone. Since levels of methionine, its two oxidation products, as well as other oxidized intermediates were higher in EC compared to VP, one could suspect elevated oxidative stress in EC. Methionine is, however, also the start of the methionine cycle generating homocysteine that is further used in the transsulfuration pathway. Finally, this pathway leads to *de novo* glutathione synthesis from methionine to regulate redox homeostasis (Fig. 4A)(*11*). In our study, certain metabolites that are part of methionine cycle (SAH) or glutathione recycling (cysteine, glycine, cys-gly, 5-oxoproline) were increased in EC compared to VP suggesting that EC exhibit greater antioxidant defense activity than VP. Taurine, that derives from cysteine via the intermediate hypotaurine, have also been reported to have antioxidant properties(*12*). Both hypotaurine and taurine levels were higher in EC compared to VP. HIV-1 infection is associated with increased ROS production and oxidative stress combined with glutathione deficiency and suppression of other antioxidant pathways(*13*). In our EC cohort though, levels of biochemicals that are part of antioxidant defense pathways were similar to the ones in HC cohort. SAH and 5-oxoproline, both used for *de novo* glutathione synthesis, had even higher levels in EC compared to HC. Cysteinylglycine was the only metabolite of the glutathione metabolism with significantly lower levels in EC compared to HC, but still higher compared to VP. This signature could reflect that EC, but not VP, are able to keep redox homeostasis at levels seen in HIV-uninfected individuals. Our finding is consistent with a recent study, where spontaneous loss of virologic control and transition from EC status was associated with increased oxidative stress and deregulated mitochondrial function prior to loss of control. Hence, oxidative stress could also be a potential biomarker of EC status(*14*). Oxidative stress might be beneficial for HIV-1 replication and glutathione deficiency is associated with impaired survival in HIV-1 infection. In contrast, glutathione treatment prevents infection of new cells through impaired virion budding and release *ex vivo*(*13, 15*).

Inflammation, ROS production, and antioxidant depletion are related with each other: Inflammatory cells release ROS at sites of inflammation leading to an imbalance between oxidants and antioxidants in favor of the oxidants and subsequently to disruption of redox signaling, consequently resulting in oxidative stress.; and without an efficient antioxidant response ROS or reactive nitrogen species (RNS) can activate intracellular signaling pathways that result in expression of pro-inflammatory genes(*16*). During ART, residual levels of viral replication are associated with persistent low-level inflammation, thus supporting HIV-1 reservoir replenishment and contributing to HIV-1 persistence(*17, 18*). Besides, increased oxidative stress during HIV-1 infection, rising either directly from the virus or indirectly from HIV-related inflammation, supports the activation of latently HIV-1-infected cells(*13*). Less is known about the role of inflammation in EC, although some studies, including an earlier study from our group, suggest that gene expression and/or levels of inflammatory markers are low and comparable to HIV-uninfected individuals(*3, 19, 20*). In contrast, one study reported increased levels of inflammatory markers in EC(*4*). This observation could be due to the heterogeneous nature of the EC.

We observed that the metabolites that are associated with inflammation reduced in EC, namely along the kynurenine pathway of tryptophan metabolism (KP), where tryptophan is metabolized to kynurenine, in the liver by tryptophan dioxygenase (TDO) and in extra-hepatic tissues by the enzyme indoleamine 2,3-dioxygenase (IDO). The activity of the latter is induced by inflammatory cytokines, for example interferon-gamma (IFNγ) and tumor necrosis factor alpha (TNFα) (Fig. 5A) (*5*). Both kynurenine and its derivative and NAD precursor quinolinate displayed significant decreases in EC relative to VP while VP showed trending (kynurenine) or significant (quinolinate) increases in these same molecules relative to HC (Fig. 5B). At the same time, levels of serotonine and tryptophan were significantly lower in VP compared to EC and HC. These data suggest that VP exhibit increased metabolism of tryptophan to kynurenine, possibly due to inflammation. This observation matches with characteristically elevated inflammation markers in treatment-naïve HIV-1-infected individuals(*21*). Relatively to VP, decreased activity of KP is observed in EC and that could be due to maintenance of physiological levels of inflammation and viral suppression in EC. We furthermore measured plasma levels of 83 soluble inflammation markers and most of them were not statistically different between EC and HC, indicating a physiological state of inflammation in EC, too. That also applies to neopterin, which is not only a pro-inflammatory marker and associated with oxidative stress, but it has been described as a diagnostic and prognostic marker in HIV-1 infection. Neopterin levels correlate positively with viral load, decrease with ART and predict HIV-1-related mortality(*22, 23*).

Interestingly, only levels of three cytokines (CCL4, CCL7, and CCL20) as well as NOS3 levels were higher in EC compared to HC. Not all of these proteins belong to classical inflammatory markers though and for the three cytokines some antiviral activities have been reported. For CCL4 and CCL7 a similar trend was also seen in our earlier study with more EC samples, although it was not statistically significant, revealing the heterogeneous characteristics of EC(*3*). CCL4 and CCL7 both bind to HIV co-receptor CCR5 with CCL4 being a known HIV-1-suppressive factor (for CCR5-tropic strains) by acting as a competitor to viral binding site(*24*). Our finding of increased CCL4 levels in EC is consistent with a study by Walker *et al*. (*25*). CCL20 was shown to have antiviral activity against HIV-1 in the female reproductive tract with a direct interaction of this chemokine with HIV-1 as the proposed mechanism of inhibition(*26*). Thus, elevated plasma levels in EC might contribute to virus control by acting as a direct antiviral(*6*).

In our earlier transcriptomics study, we observed decreased RNA levels of CCR5 in EC compared to HC in the same population(*3*). To our surprise, we did not see significant changes in the number of cells with surface expression of CCR5. Instead, we observed a lower frequency of CCR6^+^ cells in all the subpopulations investigated in EC accompanied by reduced CCR6 surface levels on both CD4^+^ and CD8^+^ T cells. This finding is in accordance with a study by Gosselin *et al*. that reported diminished frequencies of CCR6^+^ T cells in HIV-1-infected subjects, both treatment-naïve and on ART. Thus, reduction of CCR6^+^ T cells might be a consequence of HIV-1 infection, also happening in EC. The same study further revealed that, although reduced in frequency, the CCR6^+^ T cells harbored higher amounts of integrated HIV-1 DNA compared to CCR6^-^ T cells. CCR4^+^CCR6^+^ and CXCR3^+^CCR6^+^ T cells were furthermore highly permissive to HIV-1 replication, regardless of virus tropism (CCR5 or CXCR4), tested both in a cell line and in primary T cells(*27*). The CCL20/CCR6 axis has earlier been discussed in the context of HIV-1 infection. Damaged epithelial surfaces at sites of HIV-1 infection lead to release of CCL20 having anti-HIV-1 properties. CCL20 attracts CCR6^+^ cells like DCs and CD4^+^ T_H_17 cells that migrate along the CCL20 gradient to the site of infection. These cells might get infected and disseminate infection further to lymph nodes, where even more CD4^+^ cells can be infected. Since CCR6^+^ were seen to be highly permissive for HIV-1 replication and harbor HIV-1 DNA, they might particularly support HIV-1 persistence and dissemination. Thus, CCL20/CCR6 is considered to be a “double-edged sword” in regard to HIV-1 infection(*6*). In our study, CCL20 plasma levels were increased while CCR6 expression was decreased on circulating CD4^+^ and CD8^+^ T cells in EC (frequency and surface expression levels). Therefore, it seems like the CCL20-CCR6 interplay per se is not increased in EC. Albeit, it cannot be excluded that CCR6^+^-expressing cell frequencies in EC peripheral blood are low because they infiltrate in other tissues along CCL20 gradient, where they support virus replication, as it might be the case in untreated viral progressors(*27*). That needs further clarification.

The chemokine CCL7 can bind to CCR1 (not included in our analyses), CCR2, and CCR3, as well as (as an antagonist) to CCR5. No changes were seen in the expression of the CCR3 and CCR5 receptors. However, frequency of CCR2-expressing CD8^+^ T cells was reduced in EC compared to HC. As of today, HIV-1-related research has mainly focused on CCR2 expression on monocytes, including a study that reported reduced proportions of CCR2^+^ monocytes in EC. Further, a higher proportion of intermediate monocytes was observed in EC in the same study (*28*). We did not see any of these two features in our cohort. Recently, reduced frequencies and surface levels of CCR2 on CD4^+^ T cells were described in a subset of EC samples (but not in all)(*29*). Although we did not see any difference in frequencies of CCR2^+^ CD4^+^ T cells, we did also notice a reduction of CCR2 surface expression levels on CD4^+^ T cells in EC. Furthermore, frequencies and surface levels of CCR2 on CD8^+^ T cells were diminished in EC. So far, the role of CCR2 expression on CD8^+^ T cells, especially in HIV-1 infection, remains mostly elusive.

NOS3 was the fourth plasma protein with higher levels in EC relative to HC, but it was the only one not being a chemokine. NOS3 and other nitric oxide synthases convert the amino acid L-arginine and oxygen into L-citrulline and nitric oxide (NO) in a complex oxidoreductase reaction. NOS3 can also contribute to limiting immune responses regulating inflammatory processes. *In vitro* studies reported stimulatory and inhibitory effects of hydrogen sulfide (H_2_S) on expression and activity of NOS3, with H_2_S deriving from homocysteine or cysteine, thus creating a link to the transsulfuration pathway and glutathione metabolism, as discussed earlier. It is, however, unclear whether and how H_2_S affects NOS3 *in vivo*(*30*). The biological impact of elevated NOS3 levels in EC is unclear and needs further studies. Due to its immunomodulatory functions, it could be speculated that increased levels of NOS3 contribute to reduced inflammation in EC.

Even though CCL4, CCL7, and CCL20, as well as NOS3 had higher levels in EC compared to HC in our study, it seems like these proteins rather contribute to EC control state instead of having pathophysiological inflammatory effects. In contrast, NOS3 even appears to have an inflammation-reducing function; however, its exact role is unclear. Altogether, low-level inflammation and physiological antioxidant defense levels might contribute to control viral replication and viral reservoir in EC. A central question remaining is whether EC, due to functioning antioxidant defense mechanisms exhibit lower oxidative stress compared to viremic progressors, and have therefore less inflammation, thus creating an unfavorable environment for viral replication; or vice versa whether low-level inflammation is supportive for antioxidant defense stemming HIV-1 pathophysiology. Both approaches shall be considered in HIV-1 cure strategies.

One possible link between inflammation and oxidative stress, the antioxidant defense could be the oxidative phosphorylation system. It is located in the mitochondria and consists of the electron transport chain as well as ATP synthase and is responsible for mitochondrial respiration and ATP production. Hence, it is a crucial part of the cells’ energy metabolism. Oxidative phosphorylation involves oxygen and the production of ROS that, under physiological conditions, is counteracted by antioxidant defense systems. Rising numbers of immunometabolic studies in the field of HIV-1 describe increased mitochondrial respiration in CD4^+^ T cells of HIV-1-infected persons with glutamine as a source for oxidative phosphorylation. These changes in immunometabolism are associated with increased expression of ROS and inflammatory cytokines, and immunometabolic dysfunction might further mediate the development of age-related diseases(*31, 32*). Further investigating the pathway of oxidative phosphorylation might give valuable insights into HIV-1 pathogenesis and targeting it with drugs could have a promising therapeutic potential for HIV-1-infected individuals.

Our study has a few limitations that merit comments. Due to their rare occurrence, the number of EC is relatively low. However, this is one of the largest cohorts of EC which has more than median ten years of HIV-1 positivity without any treatment. Due to the recent treatment guideline for “treat-all,” it is difficult to identify the therapy naïve EC. In order to have study groups as homogeneous as possible (age-, BMI-, and gender-matched) number of EC samples and consequently also of the other two groups were even more reduced. Also, proteomics analyses were of targeted nature and did, therefore not provide a complete picture of the plasma proteome.

To summarize, low-level inflammation and physiological antioxidant defense levels observed in our EC cohort might be important factors contributing to the control of viral replication. Exploring and unraveling the processes of inflammation, oxidative pathways, and antioxidant defense as well as their implication in HIV-1 infection is of great importance for developing new therapeutic strategies.

## Materials and Methods

### Study populations

For metabolomic profiling, plasma samples were obtained from three groups of individuals: untreated HIV-1-infected individuals with viremia (viremic progressor, VP, n=16) or without viremia (EC, n=14) and HIV-negative individuals (herein mentioned as HIV-negative controls, HC, n=12). Study groups consist of female and male individuals and have comparable gender proportions. Samples are age- and BMI-matched and are part of the InfCareHIV cohort from Karolinska University Hospital, Huddinge, Sweden. EC was defined as being infected with HIV-1 for more than a year with ≥3 consecutive viral loads (VL) <75 RNA copies/ml blood (and all previous VL<1000 RNA copies/ml) or as having a known HIV-1-infection for at least ten years with two or more VL measurements of which 90% were below 400 RNA copies/mL. Patients’ characteristics are given in Table S2).

### Metabolomics

Metabolomic profiling was performed at Metabolon, Inc. (North Carolina, USA) with non-targeted mass spectrometry (MS) analysis as described recently(*33*). Briefly, samples were first prepared with an automated MicroLab STAR system (Hamilton Company) and then analyzed by four ultra-high-performance liquid chromatography-tandem mass spectrometry according to Metabolon pipeline. Biochemical components were identified based on retention time/index (RI), mass to charge ratio (m/z), and chromatographic data (MS/MS spectral data) by comparison to the Metabolon reference library that consists of more than 3300 compounds.

### Statistical analyses of metabolomics data

Doughnut charts with detected metabolites were created in Microsoft Excel. One-way ANOVA was performed to identify biochemicals that differed notably between groups (p≤0.05). For multi-group comparison, a false discovery rate (FDR) adjusted q≤0.05 was utilized. In unsupervised analyses, volcano plots were used to visualize similarities and differences between sample groups. Further, principal component analyses (PCA) were performed to picture how individual samples differ from each other. The supervised classification technique Random Forest (RF) was applied to determine which biochemicals make the largest contribution to group classification. Based on the top 30 ranked metabolites from RF analyses that contribute to group separation, supervised hierarchical clustering analyses (HCA) were performed with gplots R package(*34*) and a heatmap was generated by Pearson distance method. Supervised PCA was executed by ggplot2 R package(*35*). Correlation analyses of metabolomics data were performed with stats package, spearman correlations between specific sets of metabolites in EC. Boxplots of selected metabolites that are included in the correlation analyses were created in R Studio.

### Network analyses

Metabolites and pathways networks were created with Cytoscape ver 3.6.1(*36*). For each metabolite, fold change and p-value and q-value from Mann-Whitney U test were added to network template file. Nodes are connected objects in the network and edges connections between nodes. Metabolites are linked to subpathways and subpathways to superpathways. Gradient color and size were applied to metabolites nodes depending on fold-change.

### Targeted plasma proteomics

Plasma samples were analyzed using proximity extension assay (PEA) as described earlier(*3*). Olink Immuno-Oncology panel was applied that includes 92 plasma proteins. In addition, enzyme-linked immunosorbent assay (ELISA) was performed for neopterin (IBL International), C-reactive protein (CRP, R&D systems), and CCL20 (R&D Systems, US) as per manufacturer’s instruction.

### Statistical analyses of proteomics data

Given data distribution, non-parametric Kruskal–Wallis H test was applied to extract proteins differing at least two groups (false discovery rate (FDR) < 0.05). Partial Least Squares Discriminant Analysis (PLS-DA) was performed using selected proteins using R package ropls. Two-sided Mann Whitney U Tests were carried out using R between two different conditions (FDR < 0.05). Identified proteins are represented as boxplots using ggplot2 R package(*35*).

### Flow cytometry

Peripheral blood mononuclear cells (PBMCs) of 14 EC and 8 HC were subjected to flow cytometry analyses. Samples were thawed, washed with flow cytometry buffer (PBS with 2% fetal bovine serum and 2mM EDTA) and stained for cell surface markers for 20 minutes at room temperature. The list of antibodies used is provided in Table S3. All stainings were complemented with Live/Dead fixable near IR dye (Invitrogen, USA). After antibody incubation, cells were washed with flow cytometry buffer and fixed with 2% paraformaldehyde for 15 minutes at room temperature. Acquisition was performed on BD FACS Symphony (BD Bioscience, USA) using lasers and filter settings as indicated for BUV395, BV421, BV510, BV711, BV786, FITC, BB700, PE-CF594, APC and Near IR respectively; 355nm UV (100mW) 379/28, 405nm violet (100mW), 450/50, 525/50 (505 LP), 710/50 (685 LP), 810/40 (770 LP), 488nm blue (200mW) 530/30 (505 LP), 710/50 (685 LP), 561nm Y/G (200mW), 610/20 (600LP) and 637nm Red (140mW), 670/30 and 780/60 (750LP). Flow cytometry analysis was performed using FlowJo 10.6.2 (TreeStar, Inc, Ashland, OR), Prism 8 (GraphPad Software Inc), and visualization of complex data using Spice(*37*). A gating strategy is provided in Fig. S8. Expression frequency and levels of selected surface markers are represented as boxplots using ggplot2 R packages. T-distributed Stochastic Neighbor Embedding (t-SNE) analysis was conducted in FlowJo 10.6.1 with 2,000 iterations, 20 perplexities and a learning rate of 1,000.

## Supplementary Materials

Fig. S1. Unsupervised PCA of all samples with all metabolites.

Fig. S2. Differential expression of the metabolomics data.

Fig. S3. Random Forest (RF) analyses showing the top 30 metabolites that contribute to separation.

Fig. S4. Network analyses of the significantly different metabolites.

Fig. S5. Methionine, transsulfuration, and glutathione metabolism.

Fig. S6. Boxplots representing plasma levels of CRP and Neopterin determined by ELISA.

Fig. S7. Additional figures of flow cytometry data analyses.

Fig. S8. Gating strategy of flow cytometry data.

Table S1. Gamma-glutamyl amino acids.

Table S2. Patients’ clinical and demographic characteristics.

Table S3. Antibodies used for flow cytometry analyses and their respective properties.

## Acknowledgments

We would like to thank the study participants, nurses, and clinicians who generously supported the study. We further thank Dr. Shuba Krishnan, Dr. Soham Gupta, and Dr. Robert van Domselaar for data discussions and support.

## Funding

The study is supported by a grant from the Swedish Research Council Establishment Grant (2017-01330) to UN. AS acknowledges the funding from the Swedish research council (2016-01675) and from ALF-Stockholm County Council no 2019.

## Author contributions

UN conceived and designed the study. MS and SSA performed the laboratory experiments. MS and FM did statistical and bioinformatical analyses. MS and UN prepared the figures. MS, SSA and SNP performed the FACS analysis AS has initiated and designed the Swedish Elite controller cohort. PN and AS recruited study subjects and provided clinical data. MS wrote the first draft of the manuscript, which was then reviewed by SSA, FM, SNP, AS, and UN. All the authors approved the final version of the manuscript.

## Competing interests

The authors declare no conflicts of interest. The funders had no role in the design of the study; in the collection, analyses, or interpretation of data; in the writing of the manuscript, or in the decision to publish the results.

## Data and materials availability

All data associated with this study are present in the paper or the Supplementary Materials.

## Notes

### Competing Interest Statement

The authors have declared no competing interest.

## References and Notes

1. A. D. Olson, L. Meyer, M. Prins, R. Thiebaut, D. Gurdasani, M. Guiguet, M.-L. Chaix, P. Amornkul, A. Babiker, M. S. Sandhu, K. Porter, C. A. S. C. A. D. E. C. i. E. for, An Evaluation of HIV Elite Controller Definitions within a Large Seroconverter Cohort Collaboration. PLOS ONE 9, e86719 (2014).

2. E. Gonzalo-Gil, U. Ikediobi, R. E. Sutton, Mechanisms of Virologic Control and Clinical Characteristics of HIV+ Elite/Viremic Controllers. Yale J Biol Med 90, 245–259 (2017).

3. W. Zhang, A. T. Ambikan, M. Sperk, R. van Domselaar, P. Nowak, K. Noyan, A. Russom, A. Sonnerborg, U. Neogi, Transcriptomics and Targeted Proteomics Analysis to Gain Insights Into the Immune-control Mechanisms of HIV-1 Infected Elite Controllers. EBioMedicine 27, 40–50 (2018).

4. J. Z. Li, K. B. Arnold, J. Lo, A.-S. Dugast, J. Plants, H. J. Ribaudo, K. Cesa, A. Heisey, D. R. Kuritzkes, D. A. Lauffenburger, G. Alter, A. Landay, S. Grinspoon, F. Pereyra, Differential levels of soluble inflammatory markers by human immunodeficiency virus controller status and demographics. Open Forum Infect Dis 2, ofu117–ofu117 (2015).

5. A. T. Hopper, B. M. Campbell, H. Kao, S. A. Pintchovski, R. G. W. Staal, in Annual Reports in Medicinal Chemistry, M. C. Desai, Ed. (Academic Press, 2012), vol. 47, pp. 37–53.

6. A. Y. S. Lee, H. Körner, CCR6/CCL20 chemokine axis in human immunodeficiency virus immunity and pathogenesis. Journal of General Virology 98, 338–344 (2017).

7. R. Aebersold, M. Mann, Mass spectrometry-based proteomics. Nature 422, 198–207 (2003).

8. A. Jain, T. Kolvekar, D. R. Nair, HIV infection and lipids. Curr Opin Cardiol 33, 429–435 (2018).

9. J. Hastings, G. Owen, A. Dekker, M. Ennis, N. Kale, V. Muthukrishnan, S. Turner, N. Swainston, P. Mendes, C. Steinbeck, ChEBI in 2016: Improved services and an expanding collection of metabolites. Nucleic Acids Res 44, D1214–1219 (2016).

10. M. G. Akimov, P. V. Dudina, E. V. Fomina-Ageeva, N. M. Gretskaya, A. A. Bosaya, E. V. Rudakova, G. F. Makhaeva, G. O. Kagarlitsky, S. A. Eremin, V. I. Tsetlin, V. V. Bezuglov, Neuroprotective and Antioxidant Activity of Arachidonoyl Choline, Its Bis-Quaternized Analogues and Other Acylcholines. Doklady Biochemistry and Biophysics 491, 93–97 (2020).

11. B. C. Lee, V. N. Gladyshev, The biological significance of methionine sulfoxide stereochemistry. Free Radic Biol Med 50, 221–227 (2011).

12. H. Ripps, W. Shen, Review: taurine: a “very essential” amino acid. Mol Vis 18, 2673–2686 (2012).

13. A. V. Ivanov, V. T. Valuev-Elliston, O. N. Ivanova, S. N. Kochetkov, E. S. Starodubova, B. Bartosch, M. G. Isaguliants, Oxidative Stress during HIV Infection: Mechanisms and Consequences. Oxidative medicine and cellular longevity 2016, 8910396 (2016).

14. L. Tarancon-Diez, E. Rodriguez-Gallego, A. Rull, J. Peraire, C. Vilades, I. Portilla, M. R. Jimenez-Leon, V. Alba, P. Herrero, M. Leal, E. Ruiz-Mateos, F. Vidal, E. i. i. t. S. A. R. Network, Immunometabolism is a key factor for the persistent spontaneous elite control of HIV-1 infection. EBioMedicine 42, 86–96 (2019).

15. L. A. Herzenberg, S. C. De Rosa, J. G. Dubs, M. Roederer, M. T. Anderson, S. W. Ela, S. C. Deresinski, L. A. Herzenberg, Glutathione deficiency is associated with impaired survival in HIV disease. Proc Natl Acad Sci U S A 94, 1967–1972 (1997).

16. S. K. Biswas, Does the Interdependence between Oxidative Stress and Inflammation Explain the Antioxidant Paradox? Oxid Med Cell Longev 2016, 5698931 (2016).

17. M. Massanella, R. Fromentin, N. Chomont, Residual inflammation and viral reservoirs: alliance against an HIV cure. Curr Opin HIV AIDS 11, 234–241 (2016).

18. S. G. Deeks, R. Tracy, D. C. Douek, Systemic effects of inflammation on health during chronic HIV infection. Immunity 39, 633–645 (2013).

19. F. H. Cortes, H. H. S. de Paula, G. Bello, M. Ribeiro-Alves, S. S. D. de Azevedo, D. G. Caetano, S. L. M. Teixeira, B. Hoagland, B. Grinsztejn, V. G. Veloso, M. L. Guimaraes, M. G. Morgado, Plasmatic Levels of IL-18, IP-10, and Activated CD8(+) T Cells Are Potential Biomarkers to Identify HIV-1 Elite Controllers With a True Functional Cure Profile. Front Immunol 9, 1576 (2018).

20. H. Hocini, H. Bonnabau, C. Lacabaratz, C. Lefebvre, P. Tisserand, E. Foucat, J. D. Lelievre, O. Lambotte, A. Saez-Cirion, P. Versmisse, R. Thiebaut, Y. Levy, HIV Controllers Have Low Inflammation Associated with a Strong HIV-Specific Immune Response in Blood. J Virol 93, (2019).

21. M. Paiardini, M. Müller-Trutwin, HIV-associated chronic immune activation. Immunological reviews 254, 78–101 (2013).

22. S. P. Gieseg, G. Baxter-Parker, A. Lindsay, Neopterin, Inflammation, and Oxidative Stress: What Could We Be Missing? Antioxidants (Basel) 7, (2018).

23. M. Eisenhut, Neopterin in Diagnosis and Monitoring of Infectious Diseases. J Biomark 2013, 196432 (2013).

24. C. Blanpain, I. Migeotte, B. Lee, J. Vakili, B. J. Doranz, C. Govaerts, G. Vassart, R. W. Doms, M. Parmentier, CCR5 binds multiple CC-chemokines: MCP-3 acts as a natural antagonist. Blood 94, 1899–1905 (1999).

25. W. E. Walker, S. Kurscheid, S. Joshi, C. A. Lopez, G. Goh, M. Choi, L. Barakat, J. Francis, A. Fisher, M. Kozal, H. Zapata, A. Shaw, R. Lifton, R. E. Sutton, E. Fikrig, Increased Levels of Macrophage Inflammatory Proteins Result in Resistance to R5-Tropic HIV-1 in a Subset of Elite Controllers. J Virol 89, 5502–5514 (2015).

26. M. Ghosh, Z. Shen, T. M. Schaefer, J. V. Fahey, P. Gupta, C. R. Wira, CCL20/MIP3alpha is a novel anti-HIV-1 molecule of the human female reproductive tract. Am J Reprod Immunol 62, 60–71 (2009).

27. A. Gosselin, P. Monteiro, N. Chomont, F. Diaz-Griffero, E. A. Said, S. Fonseca, V. Wacleche, M. El-Far, M.-R. Boulassel, J.-P. Routy, R.-P. Sekaly, P. Ancuta, Peripheral blood CCR4+CCR6+ and CXCR3+CCR6+CD4+ T cells are highly permissive to HIV-1 infection. Journal of immunology (Baltimore, Md. : 1950) 184, 1604–1616 (2010).

28. S. Krishnan, E. M. P. Wilson, V. Sheikh, A. Rupert, D. Mendoza, J. Yang, R. Lempicki, S. A. Migueles, I. Sereti, Evidence for Innate Immune System Activation in HIV Type 1– Infected Elite Controllers. The Journal of Infectious Diseases 209, 931–939 (2013).

29. E. Gonzalo-Gil, P. B. Rapuano, U. Ikediobi, R. Leibowitz, S. Mehta, A. K. Coskun, J. Z. Porterfield, T. D. Lampkin, V. C. Marconi, D. Rimland, B. D. Walker, S. Deeks, R. E. Sutton, Transcriptional down-regulation of ccr5 in a subset of HIV+ controllers and their family members. Elife 8, (2019).

30. C. Bogdan, Nitric oxide synthase in innate and adaptive immunity: an update. Trends Immunol 36, 161–178 (2015).

31. T. R. Butterfield, A. L. Landay, J. J. Anzinger, Dysfunctional Immunometabolism in HIV Infection: Contributing Factors and Implications for Age-Related Comorbid Diseases. Curr HIV/AIDS Rep, (2020).

32. J. B. Spinelli, M. C. Haigis, The multifaceted contributions of mitochondria to cellular metabolism. Nat Cell Biol 20, 745–754 (2018).

33. H. Babu, M. Sperk, A. T. Ambikan, G. Rachel, V. K. Viswanathan, S. P. Tripathy, P. Nowak, L. E. Hanna, U. Neogi, Plasma Metabolic Signature and Abnormalities in HIV-Infected Individuals on Long-Term Successful Antiretroviral Therapy. Metabolites 9, (2019).

34. G. Warnes, B. Bolker, L. Bonebakker, R. Gentleman, W. Huber, A. Liaw, T. Lumley, M. Mächler, A. Magnusson, S. Möller, gplots: Various R programming tools for plotting data. (2005), vol. 2.

35. H. Wickham, ggplot2. Elegant Graphics for Data Analysis. Use R! (Springer International Publishing, New York, 2009), pp. XVI, 260.

36. P. Shannon, A. Markiel, O. Ozier, N. S. Baliga, J. T. Wang, D. Ramage, N. Amin, B. Schwikowski, T. Ideker, Cytoscape: a software environment for integrated models of biomolecular interaction networks. Genome research 13, 2498–2504 (2003).

37. M. Roederer, J. L. Nozzi, M. C. Nason, SPICE: exploration and analysis of post-cytometric complex multivariate datasets. Cytometry A 79, 167–174 (2011).

