## supplementary for "Distinct lipid profile, low-level inflammation and increased antioxidant defense as a signature in HIV-1 elite control status"

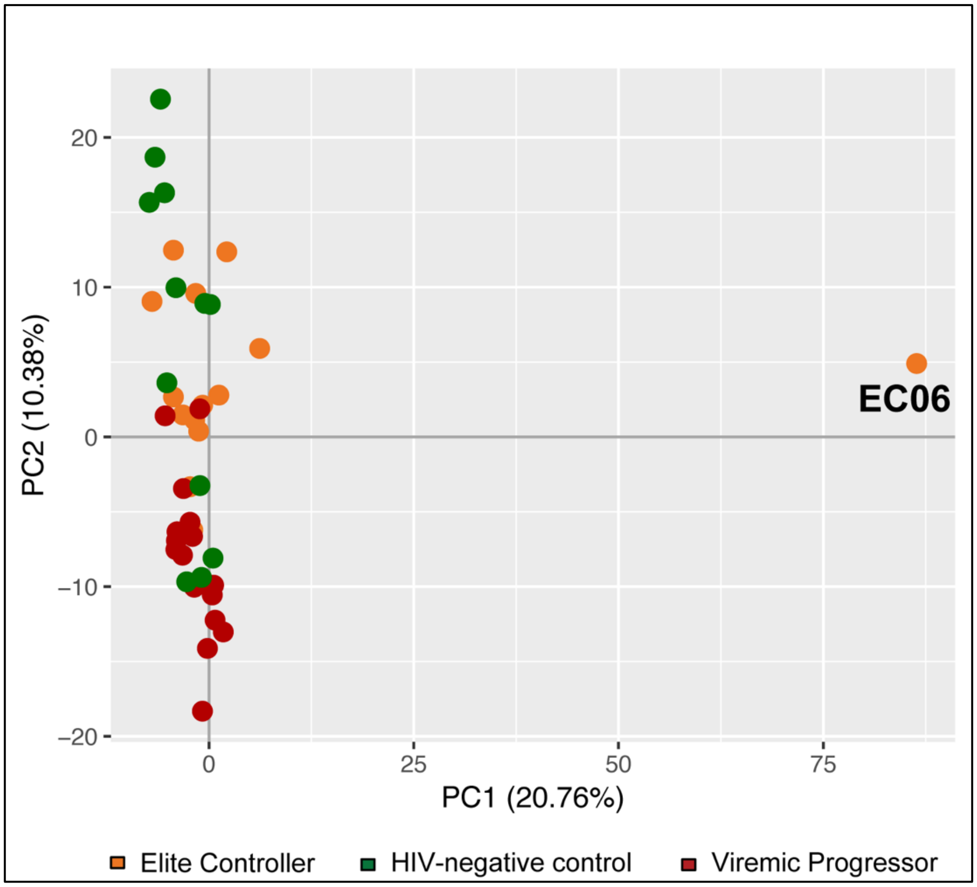


**Fig. S1.** Unsupervised PCA of all samples with all metabolites. All the samples cluster together except of EC06 that was well-separated and is there classified as an outlier. Elite controller are marked orange, HIV-negative control green, and viremic progressors red.

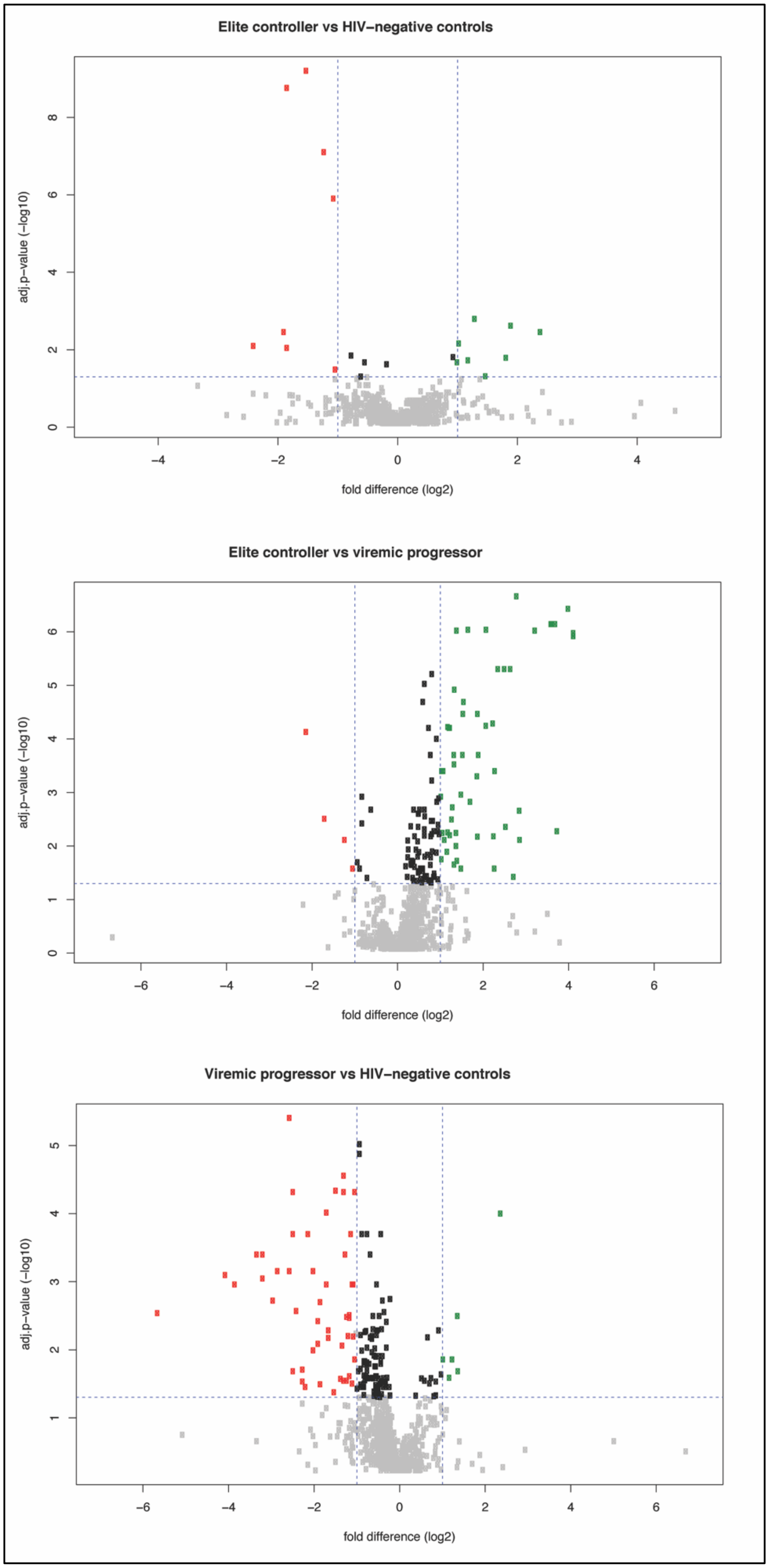

Elite controller vs HIV−negative controls
adj.p−value (−log10)
fold difference (log2)
Elite controller vs viremic progressor
adj.p−value (−log10)
fold difference (log2)
Viremic progressor vs HIV−negative controls
adj.p−value (−log10)
fold difference (log2)

**Fig. S2.** Differential expression of the metabolomics data is presented as volcano plots for EC vs HC (upper plot), EC vs VP (middle plot), and VP vs HC (lower plot). Filled red circles are used for metabolites with log2-foldchange smaller than -1.0 and adjusted p-value<0.05 (equals adj. p-value greater than $-\log_{10}1.3$). Filled green circles picture metabolites with log2-foldchange greater than 1.0 and adjusted p-value<0.05. Black filled circles constitute metabolites with log2-foldchange smaller than 1.0 and adjusted p-value<0.05. Grey filled circles are metabolites with adjusted p-value>0.05.

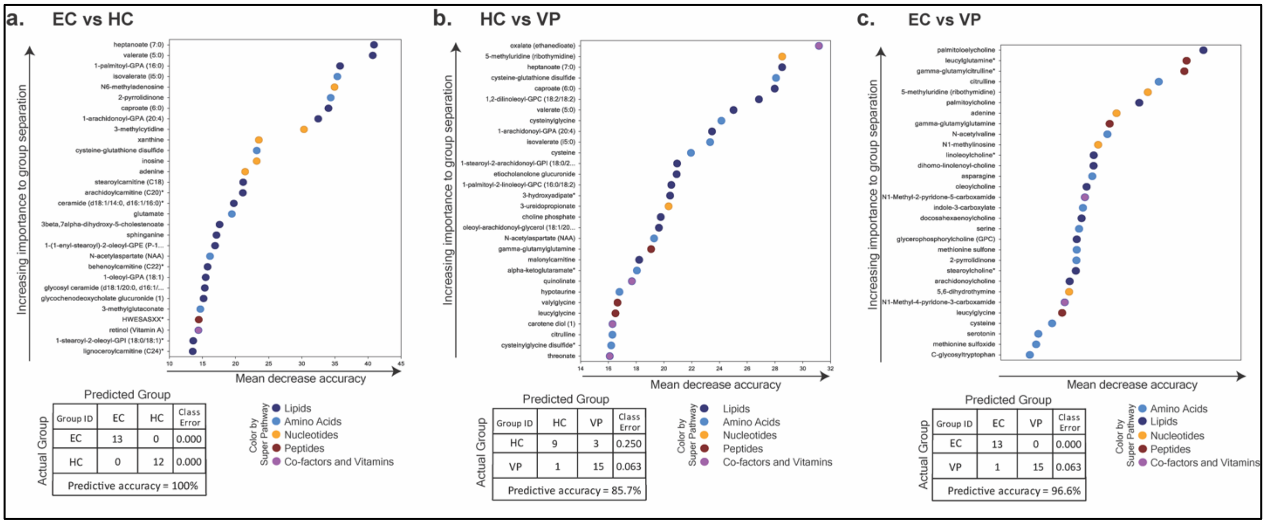


**Fig. S3.** Random Forest (RF) analyses showing the top 30 metabolites that contribute to separation between **a)** EC and HC, **b)** HC and VP, as well as **c)** EC and VP. Affiliation of each metabolite to a certain super pathway is marked by different colours (see colour-coded legend). Biochemicals involved in lipid and amino acids metabolism dominate the top-ranked intermediates for all three comparisons. The table represents predictive accuracy, predicted and actual grouping of the samples.

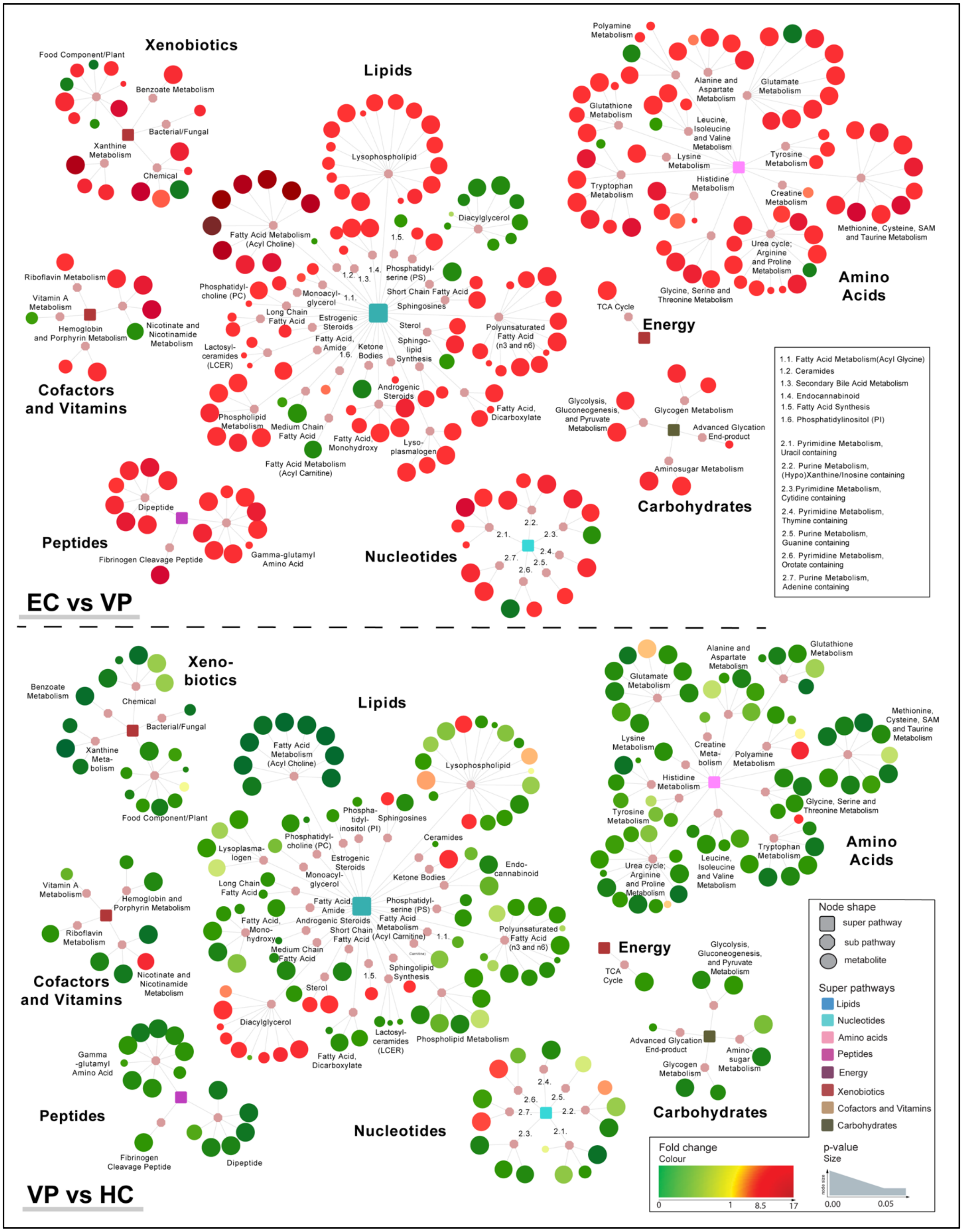


**Fig. S4.** Network analyses of the metabolites that were significantly different in EC vs VP (236 metabolites, upper network) and VP vs HC respectively (256 metabolites, lower network).

Rectangular nod shapes represent the eight super pathways that are shown in different colours according to legend. Octagonal nod shapes are used for sub pathways belonging to the eight super pathways. Circular nod shapes show the single metabolites, where red indicates increased levels and green indicates decreased levels. Size of the circles picture p-value: the bigger the size the lower the p-value. Lines connect each metabolite to its respective sub pathway and sub pathway to their respective super pathways.

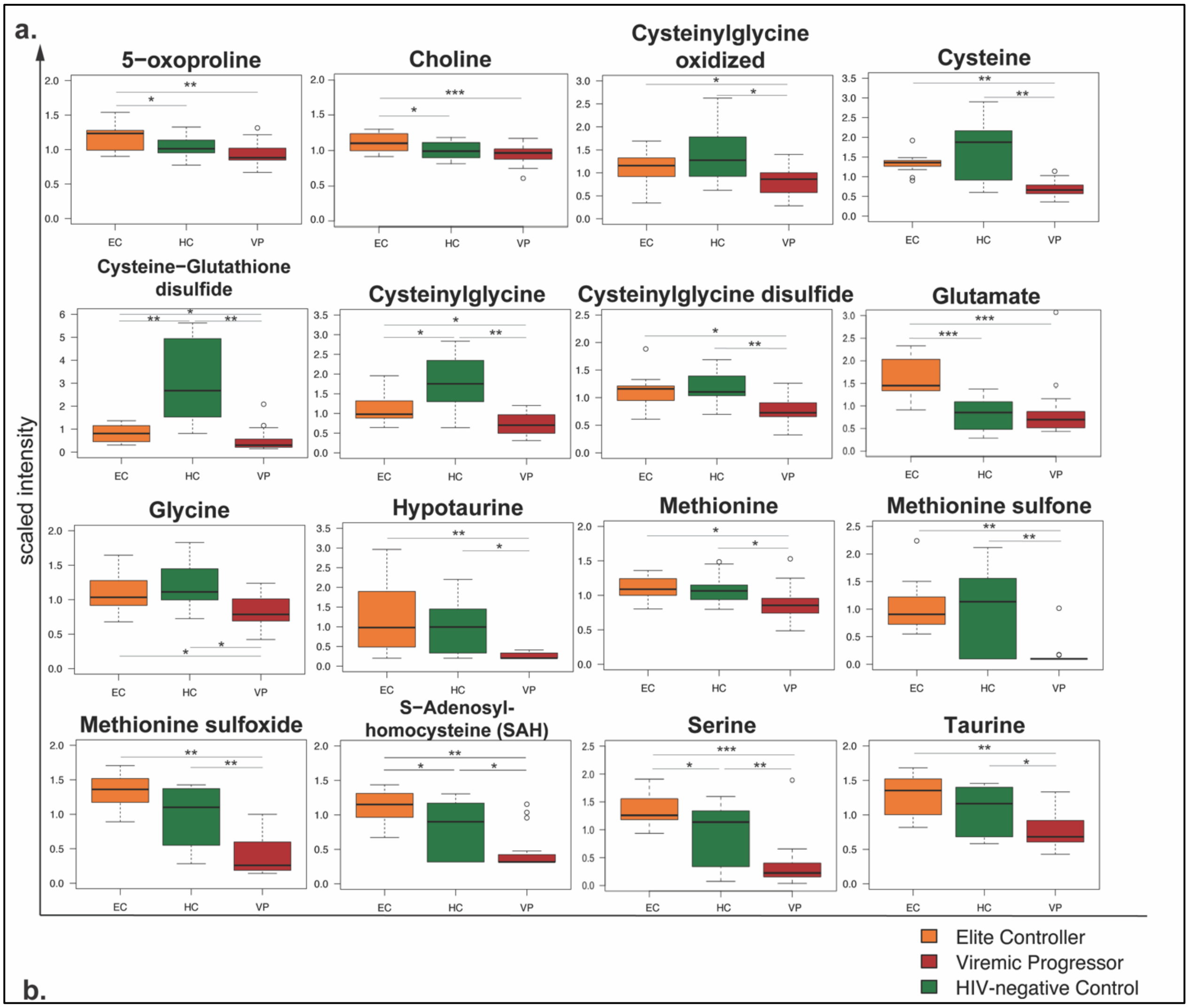


**Fig. S5.** Methionine, transsulfuration, and glutathione metabolism. Boxplots showing scaled intensity of 16 selected metabolites that play a role in methionine, transsulfuration, or glutathione metabolism. For all selected biochemicals, EC (orange) have higher levels relative to VP (red); and HC (green) has higher levels than VP for almost all metabolites shown, based on ANOVA contrasts with *indicating p-value<0.05, **p-value<0.01, and ***p-value<0.001 respectively. Median values and interquartile ranges are indicated by bars.

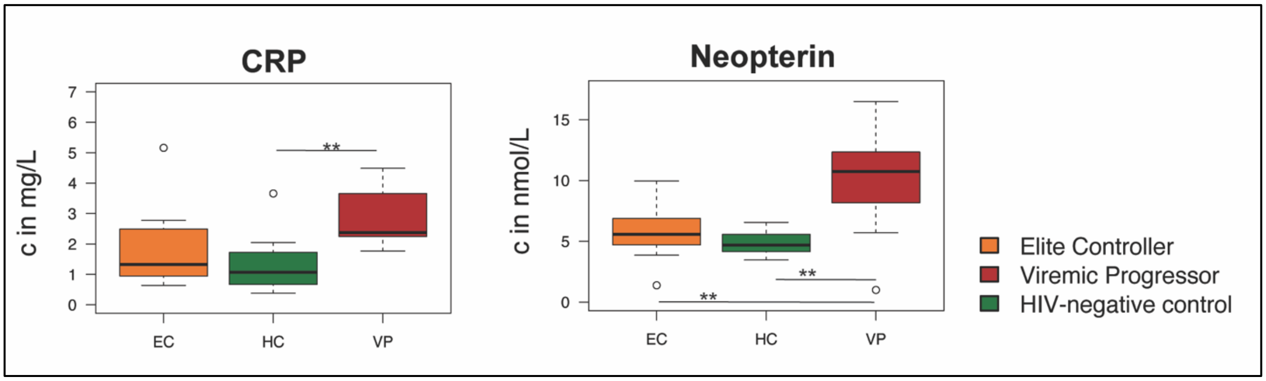


**Fig. S6.** Boxplots representing plasma levels of CCL20, CRP, and Neopterin determined by ELISA for EC (orange), HC (green), and VP (red). Concentrations are given according to unit on y-axis. Mann-Whitney test was used to calculate p-values with *p-value<0.05, **p-value<0.01, ***p-value<0.001.

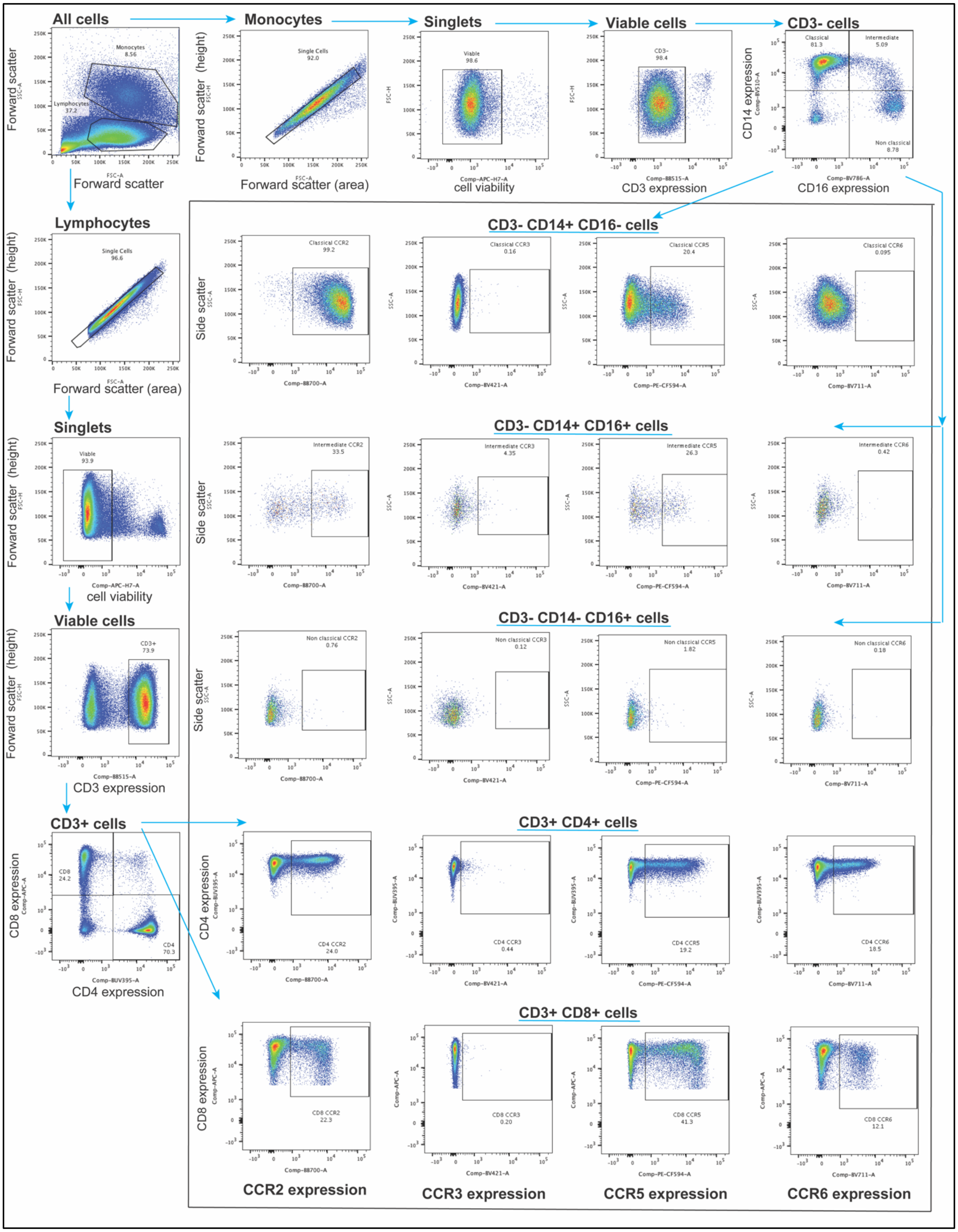


**Fig. S7.** Gating strategy of flow cytometry data.

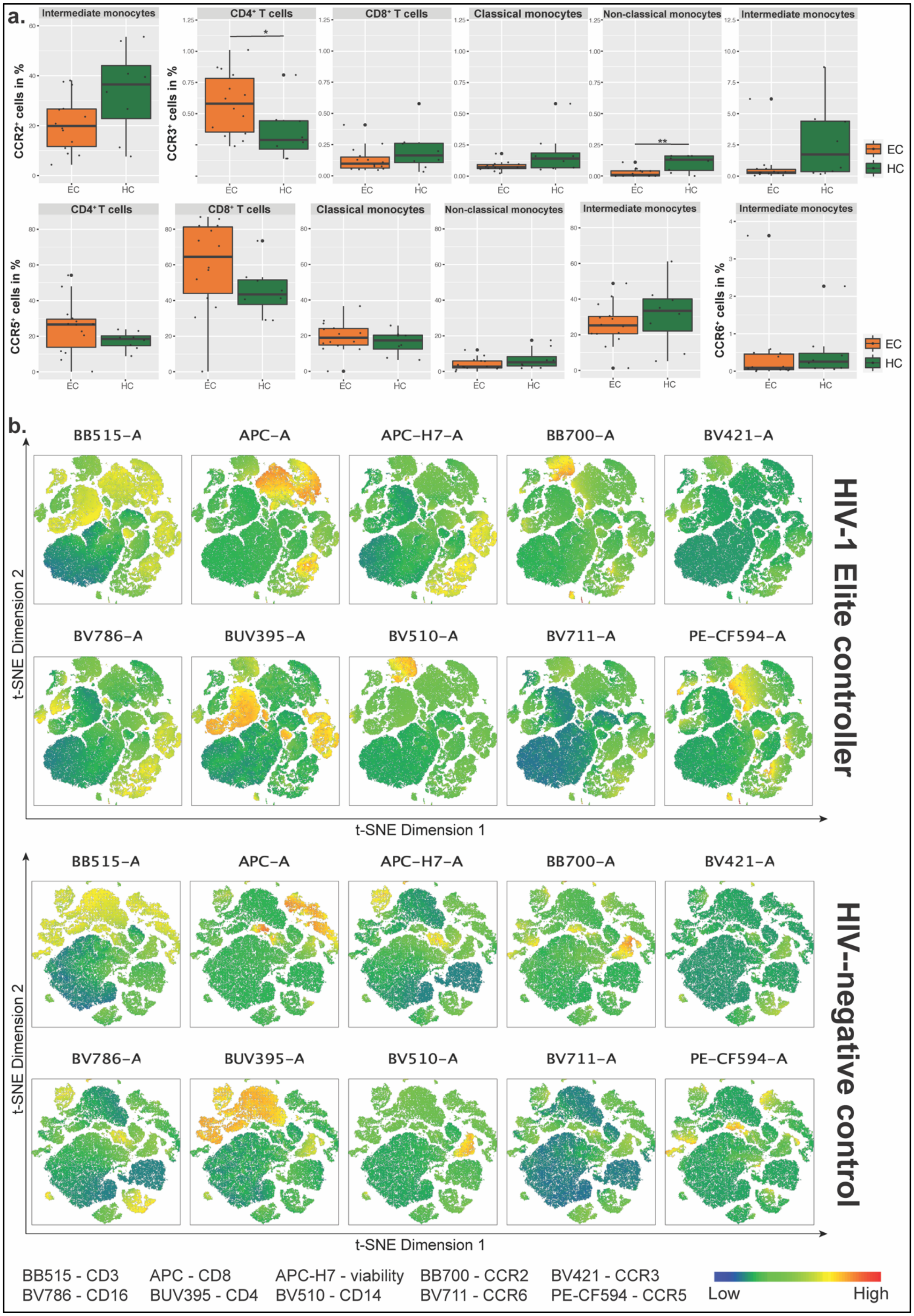
a.
Intermediate monocytes
CD4⁺ T cells
CD8⁺ T cells
Classical monocytes
Non-classical monocytes
Intermediate monocytes
CCR2⁺ cells in %
CCR3⁺ cells in %
EC
HC
CD4⁺ T cells
CD8⁺ T cells
Classical monocytes
Non-classical monocytes
Intermediate monocytes
Intermediate monocytes
CCR5⁺ cells in %
CCR6⁺ cells in %
b.
BB515-A
APC-A
APC-H7-A
BB700-A
BV421-A
BV786-A
BUV395-A
BV510-A
BV711-A
PE-CF594-A
t-SNE Dimension 2
t-SNE Dimension 1
HIV-1 Elite controller
HIV-negative control
BB515 - CD3
BV786 - CD16
APC - CD8
BUV395 - CD4
APC-H7 - viability
BV510 - CD14
BB700 - CCR2
BV711 - CCR6
BV421 - CCR3
PE-CF594 - CCR5
Low
High

**Fig. S8.** Additional figures of flow cytometry data analyses. **a)** Boxplots represent expression frequency of selected surface receptors on different cell populations for EC (orange) and HC (green). Median values and interquartile ranges are indicated by bars. P-values are determined by Mann-Whitney U test with * indicating p-value<0.05 and **p-value<0.01. **b)** t-SNE plots illustrating differences between EC (upper plot) and HC (lower plot) in surface expression of the nine targeted surface markers CD3, CD4, CD8, CD14, CD16, CCR2, CCR3, CCR5, and CCR6 respectively. The plots show t-SNE dimension 1 and t-SNE dimension 2. Blue colour shows low expression, green intermediate expression, and red high expression. Prior analysis, viable singlet cells of each sample were downsampled to 10,000 and individual downsampled samples of EC and HC respectively were concatenated. t-SNE analysis was performed with 2,000 iterations with perplexity of 20 and learning rate of 1,000.

| | | EC<br>HC | VP<br>HC | EC<br>VP | EC / HC | | VP / HC | | EC / VP | |
|---|---|---|---|---|---|---|---|---|---|---|
| | | | | | *p*-value | *q*-value | *p*-value | *q*-value | *p*-value | *q*-value |
| **Gamma-glutamyl Amino Acid** | gamma-glutamylalanine | 0,74 | 0,93 | 0,80 | 0,8396 | 0,7597 | 0,2943 | 0,3069 | 0,1994 | 0,4138 |
| | gamma-glutamylglutamate | 1,44 | 0,72 | 1,99 | 0,1498 | 0,4043 | 0,0405 | 0,0982 | 0,0006 | 0,0060 |
| | gamma-glutamylglutamine | 1,14 | 0,36 | 3,18 | 0,3049 | 0,5370 | 0,0000 | 0,0000 | 0,0000 | 0,0000 |
| | gamma-glutamylglycine | 1,09 | 0,60 | 1,83 | 0,5986 | 0,6679 | 0,0029 | 0,0181 | 0,0005 | 0,0053 |
| | gamma-glutamylhistidine | 0,99 | 0,81 | 1,21 | 0,8598 | 0,7687 | 0,0675 | 0,1305 | 0,0410 | 0,1473 |
| | gamma-glutamylisoleucine | 1,08 | 0,96 | 1,12 | 0,3872 | 0,5681 | 0,9177 | 0,5780 | 0,4109 | 0,6104 |
| | gamma-glutamylleucine | 0,93 | 0,90 | 1,04 | 0,7045 | 0,7203 | 0,4278 | 0,3788 | 0,6841 | 0,7543 |
| | gamma-glutamyl-alpha-lysine | 0,96 | 0,74 | 1,30 | 0,8940 | 0,7813 | 0,0181 | 0,0566 | 0,0223 | 0,0948 |
| | gamma-glutamyl-epsilon-lysine | 1,14 | 0,96 | 1,20 | 0,3125 | 0,5391 | 0,6012 | 0,4591 | 0,1100 | 0,2925 |
| | gamma-glutamylmethionine | 1,03 | 0,65 | 1,57 | 0,6890 | 0,7111 | 0,0007 | 0,0063 | 0,0001 | 0,0021 |
| | gamma-glutamylphenylalanine | 0,93 | 0,97 | 0,96 | 0,5009 | 0,6269 | 0,6362 | 0,4763 | 0,8109 | 0,7932 |
| | gamma-glutamylthreonine | 1,14 | 0,57 | 1,98 | 0,3026 | 0,5343 | 0,0025 | 0,0164 | 0,0001 | 0,0013 |
| | gamma-glutamyltryptophan | 1,03 | 0,71 | 1,45 | 0,7795 | 0,7421 | 0,0005 | 0,0052 | 0,0002 | 0,0021 |
| | gamma-glutamyltyrosine | 1,04 | 0,99 | 1,05 | 0,8064 | 0,7513 | 0,7870 | 0,5288 | 0,5904 | 0,7162 |
| | gamma-glutamylvaline | 1,04 | 1,08 | 0,97 | 0,7799 | 0,7421 | 0,3307 | 0,3256 | 0,4846 | 0,6664 |
| | gamma-glutamylcitrulline | 1,32 | 0,31 | 4,26 | 0,1526 | 0,4060 | 0,0000 | 0,0001 | 0,0000 | 0,0000 |
| | gamma-glutamyl-2-aminobutyrate | 1,05 | 0,83 | 1,27 | 0,6107 | 0,6698 | 0,4100 | 0,3688 | 0,1689 | 0,3861 |

**Table S1.** Gamma-glutamyl amino acids. Table with 17 metabolites belonging to the sub pathway of gamma-glutamyl amino acids that have been detected in metabolomics analyses. Foldchange between two study groups is given in column 3-5 with (EC vs HC, VP vs HC, and EC vs VP) with respective p-value and q-value in rows 6-11. Foldchange written in yellow numbers indicates statistical significance (p-value<0.05) with green cells marking downregulation and red cells marking upregulation. ANOVA contrasts were used to calculate p-values as well as q-values.

### Table S2. Patients' clinical and demographic characteristics.

| | EC | VP | HC | p-value |
|---|---|---|---|---|
| **Sample size, n** | 14 | 16 | 12 | |
| **Age, median (IQR)** | 45.5 (40.25-50.50) | 45.0 (40.75-53.25) | 44.00 (42.00-46.50) | 0.7008 (a) |
| **Gender, female, n** | 7 (50) | 7 (43.75) | 6 (50) | |
| **BMI, median (IQR)** | 26.40 (24.80-32.23) | 26.00 (23.75-30.50) | 23.00 (21.45-25.50) | 0.0537 (a) |
| **Route of transmission / Sexual orientation, n (%)** | | | | |
| **BDPT** | 2 (14.29) | | | 0.393 (b) |
| **HET** | 8 (57.14) | 10 (62.5) | 5 (41.67) | |
| **MSM** | 3 (21.43) | 4 (25.00) | - | |
| **OTR** | 1 (7.14) | - | - | |
| **PWID** | - | 2 (12.5) | | |
| **NA** | - | - | 7 (58.33) | |
| **HIV-1 subtype, n (%)** | | | | |
| **A** | | 2 (12.5) | | 0.5509 (b) |
| **A1** | | 1 (6.25) | | |
| **B** | 2 (14.29) | 2 (12.5) | | |
| **C** | 4 (28.57) | 3 (18.75) | | |
| **CRF** | 1 (7.14) | 2 (12.5) | | |
| **NA** | 7 (50) | 6 (37.5) | | |

### Table S3. Antibodies used for flow cytometry analyses and their respective properties.

| Cell surface target | Clone | Conjugate | Supplier |
|---|---|---|---|
| **CD3** | OKT3 | FITC | BioLegend |
| **CD4** | Sk3 | BUV395 | BD Bioscience |
| **CD8** | RP1-T8 | APC | BioLegend |
| **CD14** | M5E2 | BV510 | BioLegend |
| **CD16** | 3G8 | BV786 | BD Bioscience |
| **CCR2** | 1D9 | BB700 | BD Bioscience |
| **CCR3** | 5E8 | BV421 | BioLegend |
| **CCR5** | 2D7 | PE-CF594 | BD Bioscience |
| **CCR6** | G034E3 | BV711 | BioLegend |
